# Combined thermodynamic and time-resolved structural analysis of interactions between AP2 and biomimetic plasma membranes provides insights into clathrin-mediated endocytosis

**DOI:** 10.1101/2024.04.19.590255

**Authors:** Armando Maestro, Nathan R. Zaccai, J.F. Gonzalez-Martinez, P. Sanchez-Puga, J. Tajuelo, M. A. Rubio, Andreas Santamaria, J. Carrascosa-Tejedor, D. Pereira, I. Marín-Montesinos, P. Gutfreund, Richard Campbell, J. Kotar, Bernard T. Kelly, Pietro Cicuta, David J. Owen

## Abstract

Clathrin mediated endocytosis (CME) is the main mechanism for swift and selective uptake of proteins into eukaryotic cells. CME is initiated by recruitment to the plasma membrane (PM) of the adaptor protein AP2, which recognizes the PM-associated lipid PtdIns(4,5)P2, as well as the protein cargo to be internalized. Nonetheless, many aspects of this process remain unclear due to their in vivo complexity. Here, a thermodynamic and time-resolved structural analysis of AP2 binding to different biomimetic PM was undertaken under physiological conditions using a combination of neutron reflectometry, interfacial tensiometry and rheology, and atomic force microscopy. The resultant in vitro data replicated previous in vivo observations, as well as yielded biophysical insights into normal and aborted CME. The presence of cargo may not be pivotal for the “activating” conformational change of AP2. However, the presence of cargo extends AP2’s residence time on the membrane surface, due to slower on- and off-rates, thereby tentatively giving sufficient time for CME to proceed fully. Moreover, upon interaction with AP2, phospholipid lateral diffusion decreases markedly, inducing a gel phase attributed to creating a percolated network involving AP2 on the membrane, which could potentially serve as a mechanism for modulating subsequent clathrin binding.

## Introduction

Clathrin-mediated endocytosis (CME) is the primary route for swift and selective uptake of proteins into the intracellular endosomal system of eukaryotes^1^. It is crucial to controlling the plasma membrane (PM) proteome and thus plays an essential role in defining the trans-membrane protein complement of a mammalian cell’s limiting membranelle ^2^. It is therefore fundamental to cell nutrition, homeostasis, signalling and maintaining its interactions with the outside world including with other cells and the blood and immune systems. CME is also “highjacked” by many invading pathogens, including most viruses, for cellular entry and establishing infection ^3,4^. Although CME has served as a conceptual paradigm for other transport carrier formation processes ^5^, (reviewed in ^6^) many aspects of its initiation and control, and the molecular mechanisms behind them, still remain unclear at a fine-scale molecular level despite much effort.

In order to initiate CME, it has been postulated that the transmembrane PM proteins to be internalized, designated as “cargo”, need first to be selected by association with specific CME adaptor proteins ^7,8^, of which the most abundant is AP2, deletion of which in mice is lethal ^9^. Clathrin binding the PM-bound AP2 leads to the formation of clathrin-coated pits (CCPs) ^10^, which become, after subsequent scission, clathrin-coated vesicles (CCVs) carrying the cargo intracellularly.

The AP2 hetero-tetramer localizes to the cargo-presenting PM through binding the PM-specific phospholipid, phosphatidylinositol 4,5-bisphosphate (PtdIns(4,5)P_2_) ^11–13^. The presence of this phosphoinositide appears essential for CME initiation since clathrin assembly and clathrin-coated vesicle (CCV) formation are disrupted when PtdIns(4,5)P_2_ levels are lowered ^14^. Similarly, acute depletion of PtdIns(4,5)P_2_, either by a phosphatase or by inhibition of production with butanol, results in an almost complete loss of CCV formation ^15^.

Conversely, cargo over-expression leads to an increase in productive CCPs ^16^. Cargo is recognized through the binding of specific short linear amino-acid motifs present in their cytoplasmic domain. The AP2 complex recognizes two of the most commonly found internalization signals, the tyrosine YxxΦ motif (where Φ represents a hydrophobic residue) and the acidic dileucine [DE]xxxL[LI] motif ^17–20^. ‘Snap shot’ structural studies by X-ray crystallography and cryo-EM single particle reconstruction and tomography have shown that AP2 can adopt different quaternary conformations. In a closed (or locked) conformation, adopted in solution, binding sites for the tyrosine and dileucine recognition motifs are hidden ^21,22^. In an open conformational state, favoured when AP2 is ‘on-membrane’ and bound at multiple sites to PtdIns(4,5)P_2_, its cargo binding sites likely become exposed ^13,23^. Correspondingly, the clathrin binding site in the β2-hinge, which is occluded in the close-state conformation, is presumed rendered accessible on membrane binding based on protein:protein interaction and basic CCV reconstitution assays ^24^. Extrapolating from these ‘static’ molecular structures to *in cell* analysis of CME has proven insightful ^25,26,27^, but these studies suffer from a lack of dynamic physical description and energetic analysis of the collective mechanisms and driving forces, which couple AP2 cargo assembly to membrane processes. Such *in vitro* approach may provide insights to why almost half of nascent CCPs fail to mature into cargo-loaded CCVs and are instead aborted, as determined by live-cell microscopy ^8,16,28^.

In this work, characterization of bottom-up, minimal component *in vitro* PM model membranes, containing PtdIns(4,5)P_2_ and representative CME cargo, at room temperature and under physiological conditions, was followed by time-resolved biophysical measurements of these systems after addition of recombinant AP2. Low/high resolution specular neutron reflectometry (SNR) experiments were performed to study association kinetics and resultant quaternary structures of AP2 on different PM models. In parallel, the dynamic and equilibrium affinities of AP2 were determined by measurements of the membranes’ lateral pressure in the presence of increasing concentrations of AP2 using a Langmuir trough. The specific role of the lateral organization of lipids into domains, as well as the PtdIns(4,5)P_2_ function in promoting conformational change in AP2 necessary for clathrin binding, was probed by atomic force microscopy (AFM), coupled with molecular level fluorescence measurements. These imaging techniques highlighted the formation of 2-dimensional molecular features on the PM in the presence of AP2. Finally, mechanical properties measurements identified an AP2-induced transition from fluid-like to gel-like behaviour of the PM model membrane. The structural resistance against shear deformation emerging above a critical threshold was rationalized as a 2D percolation network in which the elastic behaviour is only due to an entropic configurational contribution of AP2 and PtdIns(4,5)P_2_.

Finally, these studies point the way as to how to investigate other cellular transport and membrane remodelling processes. Their intracellular locations, typically on organelles deep within the eukaryotic cell, mean they are not amenable to contemporary high spatial and temporal resolution *in cell* techniques.

## Results

### Biophysical characteristics of the plasma membrane can be recreated *in vitro*

The plasma membrane (PM) inner leaflet of mammalian cells is in a fluid state, which can be mimicked using a mixture of anionic and zwitterionic phospholipids. A model PM membrane can be formed at the air / aqueous buffer interface (see **Methods**), using a floating Langmuir monolayer prepared from 1,2-dioleoil-sn-glycero-3-phosphoetanolamine (DOPE), 1,2-dioleoil-sn-glycero-3-phosphocoline (DOPC), anionic 1,2-dioleoil-sn-glycero-3-phospho-L-serine (DOPS) and cholesterol, and further enriched with PtdIns(4,5)P_2_, and synthetic lipo-peptides, representing cargo. This 4,5 phosphoinositide is a key marker molecule for PM, while the TGN38 and CD4 peptidolipids, that display the tyrosine and the dileucine motif respectively, were used as archetypical CME cargo ^29^ (see **Supplementary Note 1**). All *in vitro* experiments were undertaken in physiological buffers ^24,30^.

At room temperature (T=21 ± 0.5 °C), the PM model membranes’ surface pressure (Π) – Area per molecule (A) isotherms, and the corresponding lateral compressibility, did not exhibit any phase transition, suggesting that these lipid monolayers are in a liquid expanded (LE) phase, which is considered equivalent to a lipid bilayer’s liquid-crystalline state (see **Figure S2**). Importantly, the presence of either the CD4 or the TGN38 lipo-peptides did not alter the existence of the LE phase (**Figure S2**).

The structure of the lipid monolayer orthogonal to the plane of the interface was determined by specular neutron reflectometry (SNR) analysis (see **Supplementary Note 8**). The PtdIns(4,5)P_2_-enriched monolayers could be effectively modelled as two layers (**Figure S10**). A layer containing the hydrophobic acyl chains is exposed to air, while the hydrophilic lipid headgroups layer is in contact with the aqueous buffer (**Table S7**). The neutron scattering length density (SLD) distribution yielded the thickness of the lipid headgroups layer as 10 ± 1Å (**Figure S10 C**). In the presence of cargo, the CD4 and TGN38 peptides were found to be confined to the headgroups layer (**Table S2**), with a solvent penetration into the hydrophobic layer of less than 12 % in all cases. Furthermore, the neutron data for the different membranes measured at surface pressure Π_0_ = 25 ± 0.5 mN/m) yielded a mean area per molecule similar to the value determined from their Π-A isotherms (**Figure S2**) as well as consistent with a compact lipid bilayer^31,32^.

### AP2 and PtdIns(4,5)P_2_ co-cluster under near-physiological conditions

The morphology of the AP2 interacting with free-standing, Langmuir lipid monolayers was studied by epifluorescence microscopy under near physiological buffer conditions (125mM Potassium Acetate, pH 7.4), initially identified in the reconstitution of AP2-mediated clathrin polymerization ^24^. In a dedicated Langmuir trough (See **Methods** and **Supplementary Note 3**), Alexa Fluor 488-labelled AP2 and TGN38-enriched PM model lipid monolayers doped with a trace amount of Bodipy-TMR labelled PtdIns(4,5)P_2_ were imaged (**Figure 1A, B** and **Figure S6**). In the absence of AP2, the fluorescent inositide formed submicrometer-sized clusters in the lipid monolayer, which is likely due to the magnesium cations’ millimolar concentration in the HKM buffer ^33^. In the presence of AP2, simultaneous colocalization in the lateral optical plane of PtdIns(4,5)P_2_ and AP2 demonstrates the preference binding of AP2 with PtdIns(4,5)P_2_. Since the lipid fluorophore is labelled in the acyl chains, the interaction between the lipid’s inositide headgroup and AP2 is expected to be unperturbed. As expected ^29^, in the absence of PtdIns(4,5)P_2_, no AP2 is observed membrane bound (see **Supplementary Note 6**).

**Figure 1.**
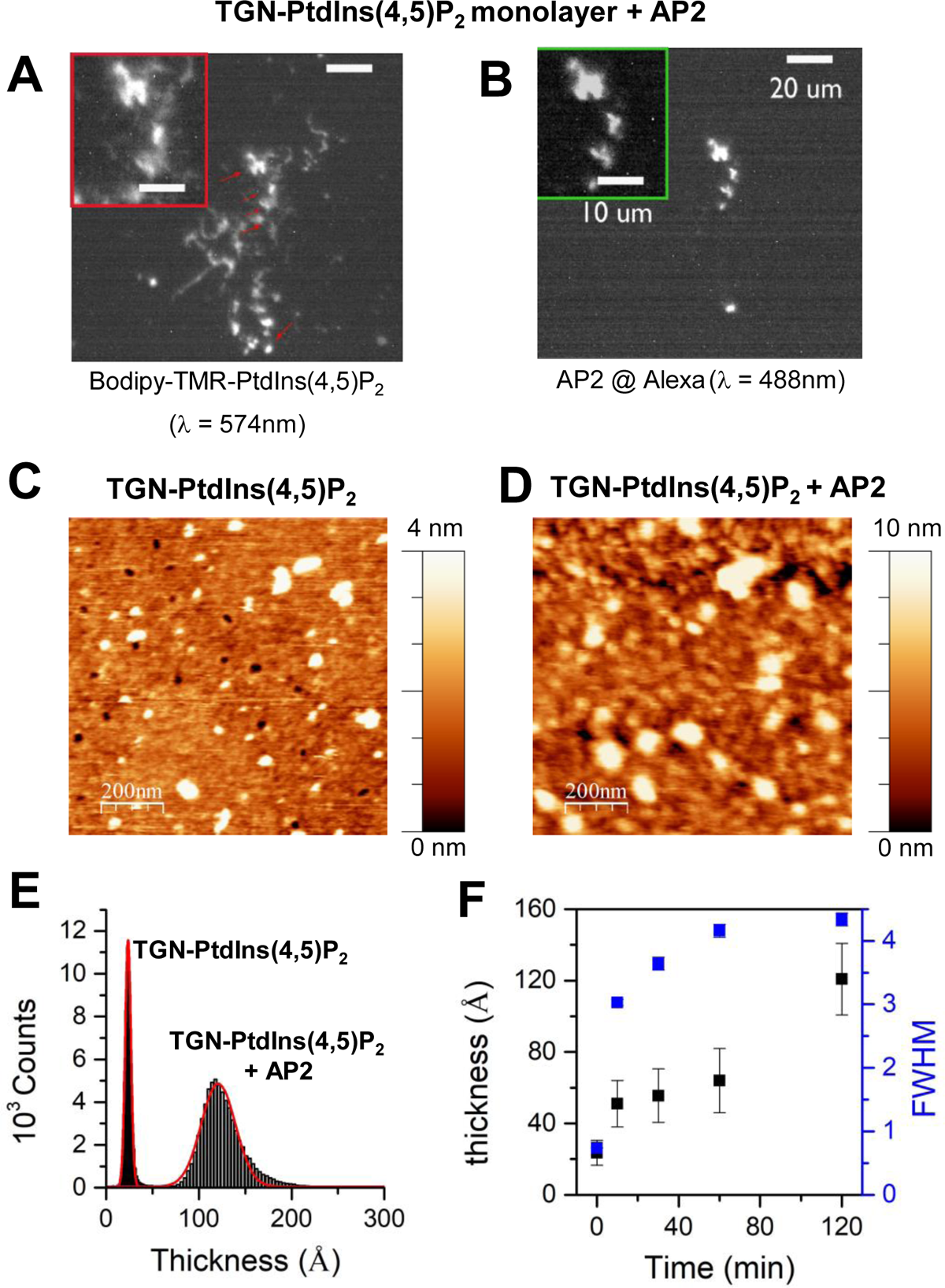
AP2 partitions into lipid monolayers enriched in PtdIns(4,5)P_2_ and TGN38: Simultaneous colocalization of AP2 and PtdIns(4,5)P_2_ clusters in Langmuir monolayers at the air water interface were reported by fluorescence images of TGN-PtdIns(4,5)P_2_ monolayer at the air/buffer interface labelled with 1 mol % Bodipy-TMR-PtdIns(4,5)P_2_ (A) interacting with AP2-labelled with Alexa Fluor 488 (B). Transferred lipid monolayers immersed in buffer were examined by fluid-phase, tapping-mode AFM in absence (C) and presence (D) of AP2. (E) shows the thickness histograms of samples shown in (C) and (D) with straight lines being the corresponding Gaussian distributions, respectively. (F) Time-resolved AFM images shows AP2 yielded an increase of thickness and FWHM after injection until a plateau is reached when AP2 saturates the monolayer. Data points are represented as mean ± S.D. (n = 10).

Tapping mode, liquid AFM investigations on Langmuir-Schaeffer supported TGN38-presenting PM model monolayers confirmed high surface coverage of the lipids, absence of defects (< 2 % surface) and the lipidic components’ miscibility (**Figure 1B**). The presence of ∼ 2nm high bright spots, which were identified as PtdIns(4,5)P_2_ clusters (≈ 8 % surface), are in agreement with previous studies of biomimetic membranes ^3435^. The corresponding monolayer lipid thickness (**Figure 1D**), is also in agreement with the above SNR analysis (30 ± 2 Å) (**Figure S10 C** and **Table S2**). Similar thicknesses were also found for CD4-enriched lipid monolayers (see **Figure S9** and **Supplementary Note 7).**

After addition of AP2, an increase in the bright spots’ size and thickness to ∼10nm is observed by AFM on TGN38-presenting PM model monolayers (**Figure 1D and 1E**). A similar increment was also found on CD4-presenting monolayers (**Figure S9**). Since in the absence of PtdIns(4,5)P_2_, such bright spots are not present (**Figure S8**), these are likely associated to AP2 binding to PtdIns(4,5)P_2_ clusters, which structural studies show occurs at 4 different places per AP2 ^13^. Time-resolved AFM shows the increase of the spot’s average thickness until a plateau is reached, which would correspond to the binding saturation of AP2 to the lipid monolayer (**Figure 1F**). Similarly, in a Langmuir trough, after injection of AP2 into the bulk buffer underneath a PtdIns(4,5)P_2_-enriched lipid monolayer, an immediate and rapid increase in lipid lateral pressure is observed, followed by a slower increase until a plateau is reached (**Figure 2A**).

**Figure 2.**
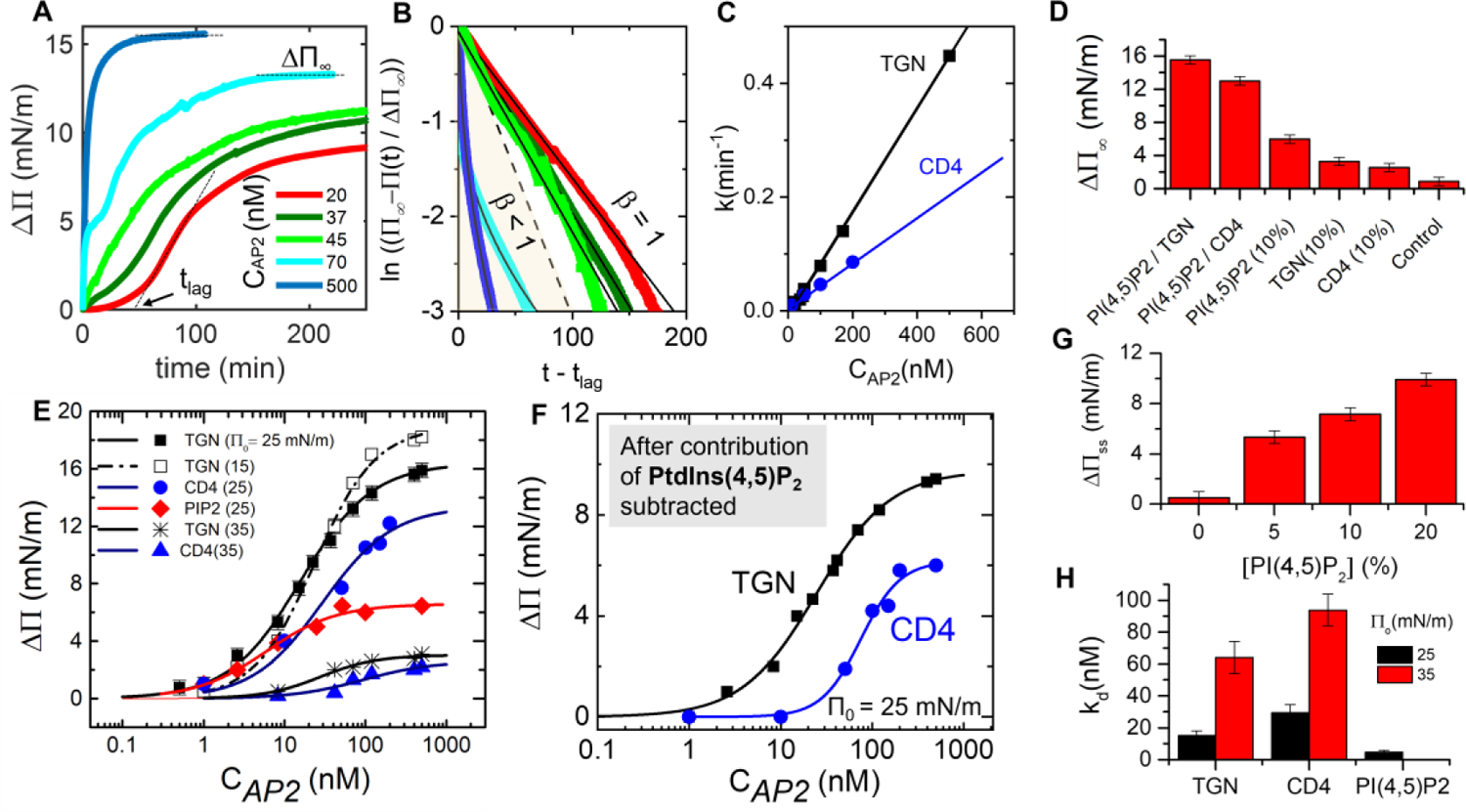
Equilibrium and kinetic tensiometry binding analysis reveal synergy between AP2 and cargo in binding PM monolayers. **(A)** Increase in Surface Pressure (ΔΠ) as a function of time for different concentration of AP2 on lipid monolayers enriched in 10 wt. % PtdIns(4,5)P_2_ and TGN38 (2wt. %). The initial surface pressure Π_0_ in all the cases has been fixed to 25 ± 1 mN/m similar to the lateral pressure in a plasma membrane leaflet. **(B)** Normalized surface pressure plotted against to the time elapsed after a lag period and fitted by a stretched exponential (straight lines) following **Equation 2. (C)** Concentration dependence of the average relaxation ratio and a linear fit (straight lines). **(D)** Maximal increase in surface pressure ΔΠ_∞_ for lipid monolayers at different composition at Π_0_ =25 ± 1 mN/m and a fixed concentration of AP2 of 500 nM. **(E)** Tensiometry binding analysis for AP2 recruited to a monolayer enriched in PtdIns(4,5)P_2_ and TGN38 or CD4 at different values of Π_0_. The increment in pressure, ΔΠ, is proportional to the amount of protein binding to the lipid monolayer at the air/water interface. Straight lines are fits to the experimental data obtained through the Hill-Langmuir relation (**Equation 1)**. **(F)** The increment in pressure, ΔΠ as a function of AP2 concentration for CD4 and TGN38 monolayers after subtracting PtdIns(4,5)P_2_ contribution. (**G**) Uppermost surface pressure increments due to the interaction of AP2 with a monolayer at Π_0_ = 25 ± 1 mN/m, at different concentration of PtdIns(4,5)P_2_. (**H**) Dissociation constants for PtdIns(4,5)P_2_ monolayers enriched in CD4 and TGn38 at different initial surface pressure. Bar diagrams in D, G, H are represented as mean ± S.D. (n = 5).

### Equilibrium binding analysis of AP2 to a model PM shows synergy between PtdIns(4,5)P_2_ and cargo binding

The binding affinity of AP2 to the lipid monolayer can be directly determined through titration of surface pressure measurements. The analysis with a Langmuir adsorption model assumes the maximum increase in the lateral pressure ΔΠ_∞_, at constant surface area, is dependent on the fraction of AP2 (θ) interacting with the monolayer, and the resultant change in lateral pressure ΔΠ^36^ (see further details on **Supplementary Note 3**). The binding affinity of AP2 to the different lipid monolayers could therefore be determined from equilibrium analysis (**Figure 2E, Table 1**),

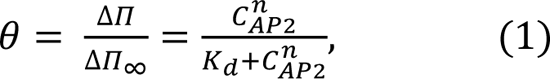

where C_AP2_ is the bulk concentration of AP2, K_d_ is the dissociation constant and n is the Hill coefficient^37^.

**Table 1:** Langmuir derived Equilibrium and kinetic binding terms for AP2 to different biomimetic PM.

| | Kinetic binding analysis | | | | Equilibrium data $\Pi = 25\text{mN/m}$ | | | |
| --- | --- | --- | --- | --- | --- | --- | --- | --- |
| | $K_{\text{on}}/10^4$<br>$\text{s}^{-1} \text{M}^{-1}$ | $K_{\text{off}}$<br>$/10^{-4} \text{s}^{-1}$ | Residence<br>time / s | $K_d$<br>/nM | $K_d$<br>/nM | $\beta$ | $K_d' (*)$<br>/nM | $\beta' (*)$ |
| PtdIns(4,5)P<sub>2</sub> | $7.9 \pm 0.7$ | $5.6 \pm 0.5$ | 1785 | $7 \pm 1$ | $5 \pm 1$ | $0.9 \pm 0.1$ | | |
| TGN+ PtdIns(4,5)P<sub>2</sub> | $1.5 \pm 0.1$ | $1.8 \pm 0.1$ | 5555 | $12 \pm 1$ | $15 \pm 3$ | $1.0 \pm 0.1$ | $25 \pm 6$ | $1.1 \pm 0.1$ |
| CD4+ PtdIns(4,5)P<sub>2</sub> | $65 \pm 2$ | 1.2 | 8333 | $18 \pm 3$ | $29 \pm 5$ | $1.1 \pm 0.1$ | $75 \pm 9$ | $1.9 \pm 0.4$ |

(\*) After contribution of PtdIns(4,5)P<sub>2</sub> subtracted.
$\beta > 1$ indicates a cooperative scenario according to the Hill equation.

At the typical membrane lateral pressure Π_0_ = 25 ± 0.5 mN/m, the affinity of AP2 to a 5% PtdIns(4,5)P_2_ PM monolayer was K_D_ = 5 ± 1 nM. The increase in concentration of PtdIns(4,5)P_2_ leads to an increase in ΔΠ_∞_ (**Figure 2G, S3**) and can be rationalized by an increased in AP2 binding capacity of the membrane (as observed in **Figure 1A**).

The addition of either TGN38 or CD4 cargo also yield further increases in lateral pressure ΔΠ_∞_, reflecting the synergy in binding AP2 from inositol phospholipids and from cargo present in the membrane (**Figure 2C, S3)**. The cargo binding curves at low AP2 concentration mirror the binding curve for PtdIns(4,5)P_2_-only monolayer, but as AP2 concentration increased, these curves reached higher equilibrium pressure changes. This biphasic binding behaviour is observed at AP2 concentrations above 2 nM for TGN38, and 10 nM for CD4. Assuming independent binding sites for PtdIns(4,5)P_2_ and for cargo on AP2, the removal of the PtdIns(4,5)P_2_ binding effect results in an equilibrium-derived K_D_ = 24.5 ± 1.6 nM and 75 ± 9 nM, for TGN38 and for CD4 respectively (**Figure 2F, Table 1**). The lower affinity for TGN38 peptide to its binding site on AP2 determined in solution by ITC (2 μM) suggest lipid head groups also contribute to AP2 binding^25^. In support of this observation, AP2 has an overall weaker affinity to more densely packed PM model monolayers, which had larger initial pressure Π_0_ = 35 ± 0.5 mN/m (overall affinity weakening from K_D_ = 15 ± 3 nM to 64 ± 10 nM and from 29.5 ± 5 nM to 93 ± 10 nM, for a TGN- and a CD4-enriched PM monolayer respectively) as shown **Figure 2G**.

These affinities between AP2 and model PM appear to mirror the relative difference in affinities previously reported from SPR Biacore measurements on immobilized liposomes ^11^. Although, the SPR-derived affinities were significantly weaker (311 nM for PtdIns(4,5)P_2_ and TGN-enriched liposomes, and 930 nM for PtdIns(4,5)P_2_ and CD4-enriched liposomes), this could be due to the higher salt concentration in the SPR buffer (250mM NaCl), which would reduce any ionic interaction between PtdIns(4,5)P_2_ and AP2, in comparison to the more physiological buffer used (125 mM Potassium Acetate).

### Kinetic binding analysis shows that the presence of cargo significantly increases AP2 residence time on model PM

The binding affinity obtained by the Langmuir isotherms at equilibrium (steady state) were complemented by kinetic experiments to evaluate the temporal behaviour of the association/dissociation of the AP2-lipid monolayer binding process (**Figure 2 A, B)**. Under 70nM AP2 solution concentration, the kinetics display an exponential trend, while for concentrations above 70nM, the kinetics can be fitted by a stretched exponential expression based on rate distributions ^38^ (see further details in **Supplementary Note 3**),

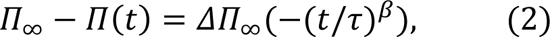

which are determined by the characteristic time constant τ and the stretched exponent β (0 ˂ β ˂ 1). The latter parameter controls the width of the corresponding rate distribution. At increasing AP2 concentrations, a decrease in both τ and β are observed (See **Table S5**). An average binding time ˂τ˃ was calculated from

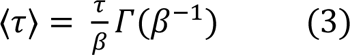

Where Γ denotes the gamma function ^39^. The associated relaxation rate k_d_ = τ^-1^, increases linearly with AP2 concentration (**Figure 2D**). Assuming a first order kinetic reaction (see SI Methods), the apparent rate coefficient is given by

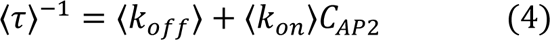

Where the apparent k_on_ on rate and k_off_ off rate take into account the combined effect of cargo and PtdIns(4,5)P_2_ binding.

The intercept and slope of a linear fit to the experimental values of ˂τ˃^-1^ (**Figure 2G**) yield the average backward and forward rate coefficients for AP2 association to PtdIns(4,5)P_2_-only membranes giving 〈*k*_*off*_〉 = 5.6 ± 0.5 10^−4^*S*^−1^ and 〈*k*_*on*_〉 = 7.9 ± 0.7 10^4^*M*^−1^*S*^−1^, respectively. At 25 mN/m, the ratio 〈*k*_*off*_〉/〈*k*_*on*_〉 gives an average dissociation coefficient K_D_ (7 ± 1 nM), which is similar to the corresponding value of (4.7 ± 0.5 nM) from the equilibrium analysis of the Langmuir isotherm (**Figure 2G)**.

In the presence of cargo, the affinities determined from both the kinetic and the equilibrium analysis of the Langmuir isotherm agree (12 ± 1 and 18 ± 3 nM in the presence of TGN38 and CD4 respectively) (**Figure 2G, Table 1)**. However, in comparison to cargo-less PM model membranes, the kinetic reaction slows down remarkably, with both the k_on_ and k_off_ rates reduced approximately three-fold (see table). As a result, the mean residence time of AP2 on the model membrane increases significantly from 1785 s (30 minutes) to 5555 s (90 minutes) and 8333 s (∼140 minutes) in the presence of TGN38 and CD4 cargo respectively, which is in line with the measured nanomolar K_D_. The cargo therefore not only stabilizes open conformation of AP2 ^11,24,40^, but also its presence on the membrane. The cargo-driven increase in residence time at the membrane surface, we believe allows sufficient clathrin to bind and therefore productive nucleation of clathrin coated pits to proceed. A similar effect was observed *in cell* by comparing wild type and mutant AP2 (the latter unable to bind specific cargo) ^23,26^.

Similar agreement was also found between K_d_ calculated by the kinetic analysis and the Langmuir isotherm for the binding of CD4 and PtdIns(4,5)P_2_ monolayers. However, the non-exponential kinetics present at high bulk concentrations of AP2 would imply a more complex reaction scheme that cannot be adequately captured by average rate coefficients. This was also suggested by the complex analysis of AP2 liposome SPR “binding curves”^29^.

### Time-resolved AP2 binding to model PM measured by SNR and by tensiometry agree

Time-resolved specular neutron reflectometry (SNR) was used to directly measure the AP2 concentration associated to model PM **(Supplementary Note 3** and **Figure S20).** By comparing the surface pressure measured in parallel, pressure increase and SNR-derived membrane bound protein concentration follow similar trends as shown **Figure 3 A**. A linear dependence ΔΠ ≈ Γ/G∞ ≈ ϕ, as shown in **Figure 3 B**, indicates that the protein recruited by the lipid monolayer directly contributes to the rise in lateral pressure. ϕ represents here the surface coverage of AP2. Normalized plots of the recruitment of AP2 to different model PM are shown in **Figure 3 G**. After an induction period t < t_lag_, there is an increasing number of AP2 recruited to the membrane. The linear fit to the SNR data corresponds to a first order binding reaction (See **Figure 3 G** and **Supplementary Note 3**). The values of binding time scale τ, reported in **Figure 3 H**, are in the same order, although slightly lower in the case of TGN38-monolayers, which were obtained from fitting the temporal evolution of the surface pressure data shown in **Figure 2 B**. Moreover, similar trends were observed by the temporal evolution of elliptically polarized light, which can be defined in terms of the ellipsometric angles that are proportional to the quantity of AP2 recruited to different PM monolayers per unit area (**Figure S5** and **Supplementary Note 5**).

**Figure 3.**
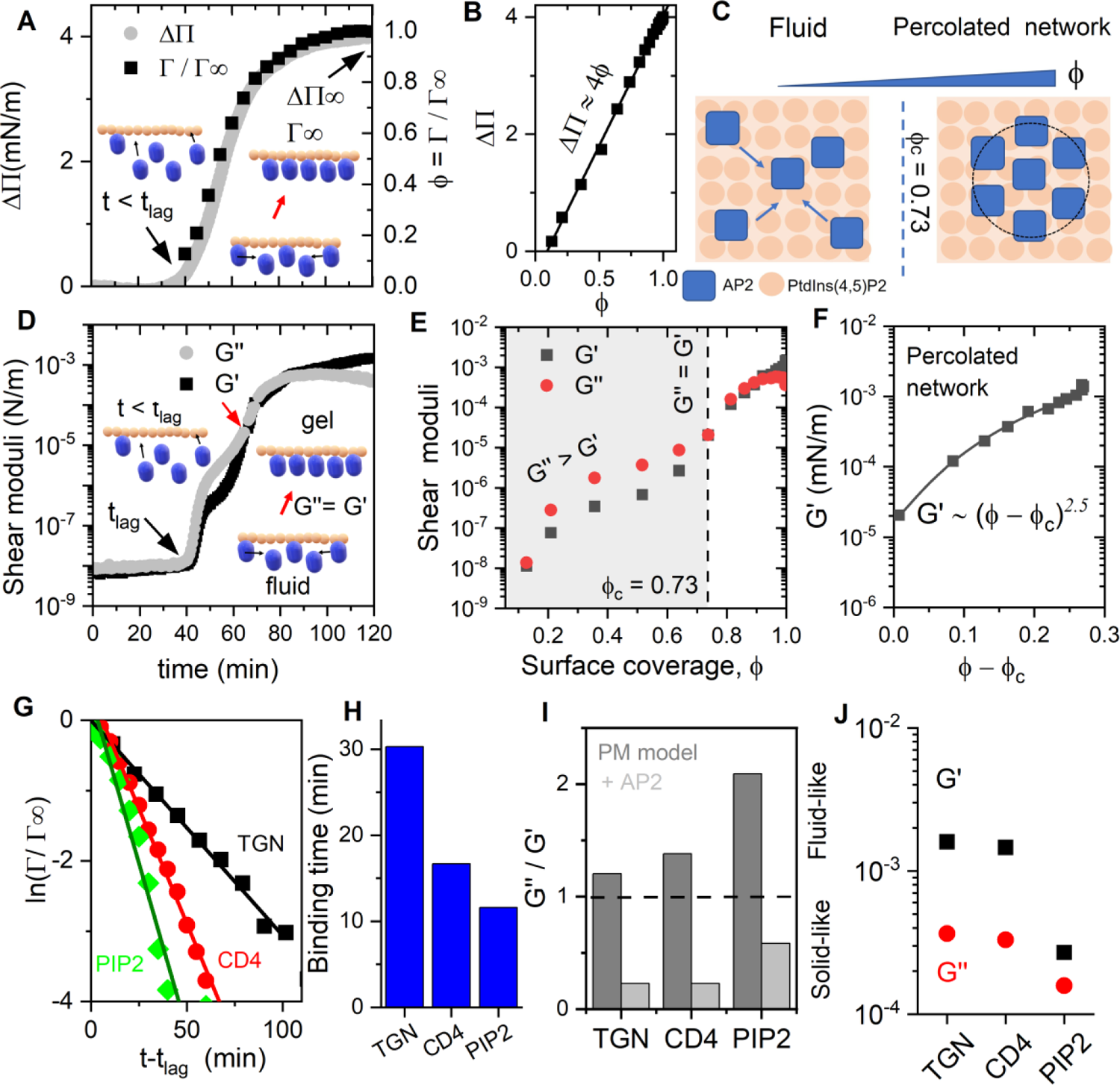
Combination of time-resolved neutron reflectometry and interfacial shear rheology shows the existence of a AP2/PtdIns(4,5)P_2_ entropic network at increasing surface coverage. (A) Normalized amount of AP2 protein (Γ/G∞, with Γ∞ being the saturation concentration) interacting with a lipid monolayer enriched in PtdIns(4,5)P_2_ and TGN38 measured using time-resolved SNR in a restricted Q region and compared with the increase in surface pressure measured (ΔΠ). (B) Linear dependence between ΔΠ and the fraction coverage (ϕ = Γ/G∞). (C) The suggested transition mechanism from a fluid state to a percolated network state occurs as the coverage of AP2 on a PtdIns(4,5)P_2_-enriched membrane (ϕ) rises. (D) Time dependence of the interfacial shear viscoelastic moduli, divided in the elastic and viscous modulus (G’ and G”, respectively). Data shows an increase in both G’ and G’’ after injecting AP2 in the subphase. (E) Re-plotted G’ and G’’ with respect to the surface coverage ϕ, revealing two differentiated regimes: a fluid-dominated state below ϕ = 0.73 and a gel-like state above this threshold that can be rationalized as percolated network according to (C). (F) G’ plotted as a function of the distance to the percolation threshold showing a power law exponent *f* = 2.3. (G) Comparative for monolayers enriched with and without different cargoes such as TGN38 and CD4. Straight lines are fits of the experimental data to obtain an average binding time plotted in H. G’’/G’ ratio of the different monolayers studied before and after the injection of AP2 is plotted in I. (J) shows the values of G’ and G” for monolayers with different composition. The initial surface pressure Π_0_ in all the cases has been fixed to 25 ± 1 mN/m and the bulk concentration of the protein to 10 nM.

### Fluid to solid-like transition in model PM monolayers on binding of AP2

In order to complement *in situ* measurements of lateral surface pressure, AP2 binding to model PM was monitored by interfacial shear rheological measurements (**Methods** and **Supplementary Note 4**). These data yielded the model PM lipid monolayer change in shear storage G’(t) and loss modulus G’’(t) (**Figure 3D** and **Figure S4**). Prior to AP2 injection, the different PtdIns(4,5)P_2_ enriched monolayers are in a fluid state, denoted by G’’/G’ ≥ 1 (**Figure 3 I, J**). On injection of AP2, after an induction period t < t_lag_, both G’ and G’’ increase ∼10,000 fold before reaching an equilibrium, at which G’’/G’ < 1 indicates an elastic response that is characteristic of a solid-like regime. Moreover, in the presence of peptide-lipid cargo, the resultant moduli are further increased, with respect to PtdIns(4,5)P_2_ only monolayers, which may be rationalized by the increased number of AP2 recruited by the cargo-presenting PM monolayer. The higher shear modulus would imply a drastic reduction in phospholipid mobility. The tightly packed lipid structure would tentatively generate strong intermolecular forces and resistance to shear deformation induced by clathrin polymerization.

The dependence of shear storage G’ and loss modulus G’’ with AP2 coverage is captured in **Figure 3E**. Importantly, a critical surface threshold ϕ_c_ = 0.73 can be visualized when G’ = G’’ and associated to a randomly close packed distribution in 2D ^41^. Beyond ϕ_c_ a disordered gel phase characterized by G’ > G’’ can be structurally hypothesized in terms of a percolation network (Stauffer and Coniglio 1982). It is assumed that isolated AP2 molecules bound to the membrane come into mutual contact and randomly arranged forming a connected network (as represented in **Figure 3C**). Therefore, G’ can be described by a power law relation above the critical percolation concentration (ϕ − ϕ_c_)^f^, with *f* = 2.3, which is typical of a protein network distribution ^43^ (**Figure 3F**). As a corollary, AP2 is able to create a solid but fragile connected network with a purely elastic modulus that is entropic in origin assuming excluded volume interactions ^44^. This percolation network approach relies solely on a statistical physics framework of connected clusters, where network formation is solely influenced by molecular packing of AP2, and not by specific details of protein recruitment and reorganization. The connected network formed by AP2 could influence subsequent clathrin lattice assembly.

### AP2 binds PM headgroups but embeds no further as determined by low-resolution SNR

Due to the sub-micromolar binding affinity of AP2 to model PM monolayers, the SNR experimental set up was adapted for full-Q_z_ range analysis to determine the structures of AP2 bound to lipid monolayers. **Figure S12** shows the reflectivity profiles measured as a function of Q_z_ in two isotropic contrasts (38% D_2_O and 100% D_2_O, see methods), after the binding of AP2 to different model PM monolayers. Data modelling was performed by simultaneously fitting the two contrasts to get a single set of structural parameters (e.g., thickness, scattering length density, roughness and volume fraction), which also include the volume fraction of protein partitioning into different regions of the lipid monolayers (see **Table S7** and **Supplementary Note 8** for further details). The shape of AP2 was initially modelled as a series of stratified layers (See scheme in **Figure 4**). The SNR data from PtdIns(4,5)P_2_ presenting monolayer, in the presence and absence of different cargo peptides (CD4 and TGN38) indicated that in each case, AP2 lies primarily outside the membrane with a little insertion at the level of lipid headgroups (≈ 5 ± 1%), and a negligible presence in the acyl chain region (**Table S8**). A similar result was obtained by the SNR analysis of solid-supported bilayers (SLBs), composed of PtdIns(4,5)P2 enriched bilayers and CD4 cargo (**Figure S21** and **Supplementary Note 9**). However, the volume fraction of protein bound to the membrane varied significantly between PM model membranes (**Figure 4A-C**). In comparison to CD4, the presence of TGN38 leads to a larger fraction of AP2 bound to the membrane (**Figure 4** and **Table S8**). The smallest surface coverage was in the absence of cargo, on PtdIns(4,5)P_2_-only membrane. These SNR data agree with the measured relative increases in membrane lateral pressure and in shear moduli (**Figures 2 and 3**).

**Figure 4.**
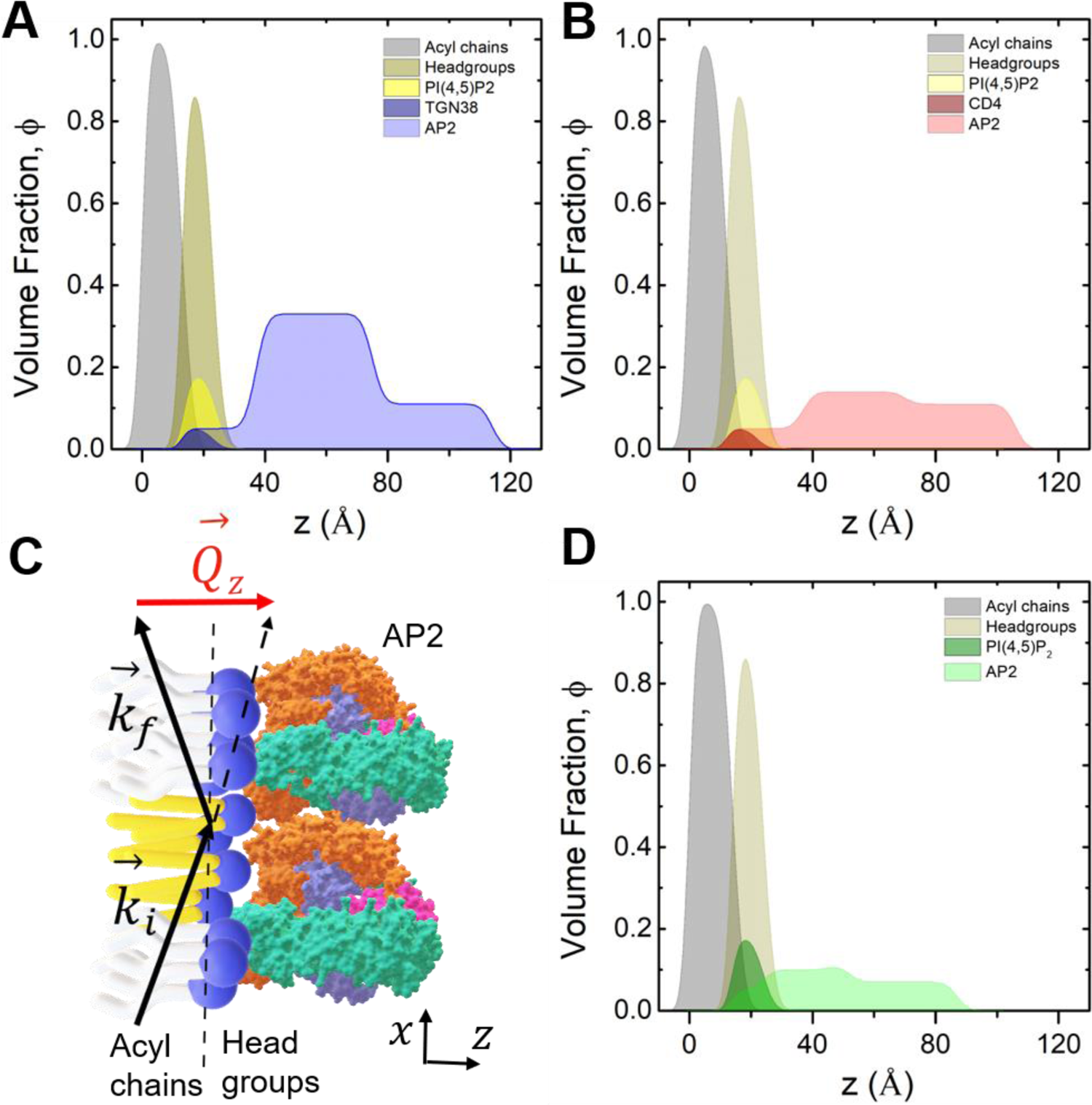
SNR analysis of AP2 binding PtdIns(4,5)P_2_ – enriched lipid monolayers shows cargo-dependent surface coverage of AP2. A, B, D shows the in-plane averaged spatial distributions profiles of TGN, CD4 and no cargo membranes respectively, expressed as volume fractions ϕ, of monolayer components and AP2 in the direction orthogonal to the plane of the interface. A scheme of the SNR profile is shown in C.

The SNR-derived low-resolution structural models of membrane-bound AP2 show in the case of TGN38 and of CD4, total protein thickness are similar (91 and 84 Å respectively) (**Table S8**). AP2 would be oriented with its major axis parallel to the monolayer normal, as the protein’s maximum width is ∼ 90 Å^40^. In the absence of cargo, the protein has reduced surface coverage and is more compact (63 Å thickness), which is in agreement with the in-plane AP2-PtdIns(4,5)P_2_ cluster formation visualized by AFM and epifluorescence microscopy (**Figure 1 A, C**).

### AP2 binds in an open conformation to model PM in the presence and absence of cargo

The initial SNR data analysis of membrane-bound AP2 yielded a low-resolution slab model of constant scattering length density (SLD) that is intrinsically linked to protein’s quaternary structure. Consequently, the preferred conformation and orientation of AP2 on the membrane can also be determined from the SNR data. In order to identify the preferred orientation and position of AP2 with respect to the PM membrane, SLD profiles were calculated from atomic models of open, open+ and closed AP2 (based on energy minimized structures from PDB 2ax7, 6QH6 and from 2vgl, respectively). An ensemble of SLD profiles from 256 different orientations for each structure were generated with 5° increments in the Euler angles α and β (following an x-y-z extrinsic rotation scheme; the third Euler angle γ is not relevant as the SLD is plane-averaged in the z-direction). After each rotation, the structure’s distance with the membrane was optimised with the Sassie software by fitting the resultant theoretical and experimental SLD^45^. In the final refinement step, the most favourable orientation for each protein conformation is fitted against the experimental reflectivity data. (See **Figure 5A, B** and **S14-S17** and **Supplementary Note 8**).

**Figure 5.**
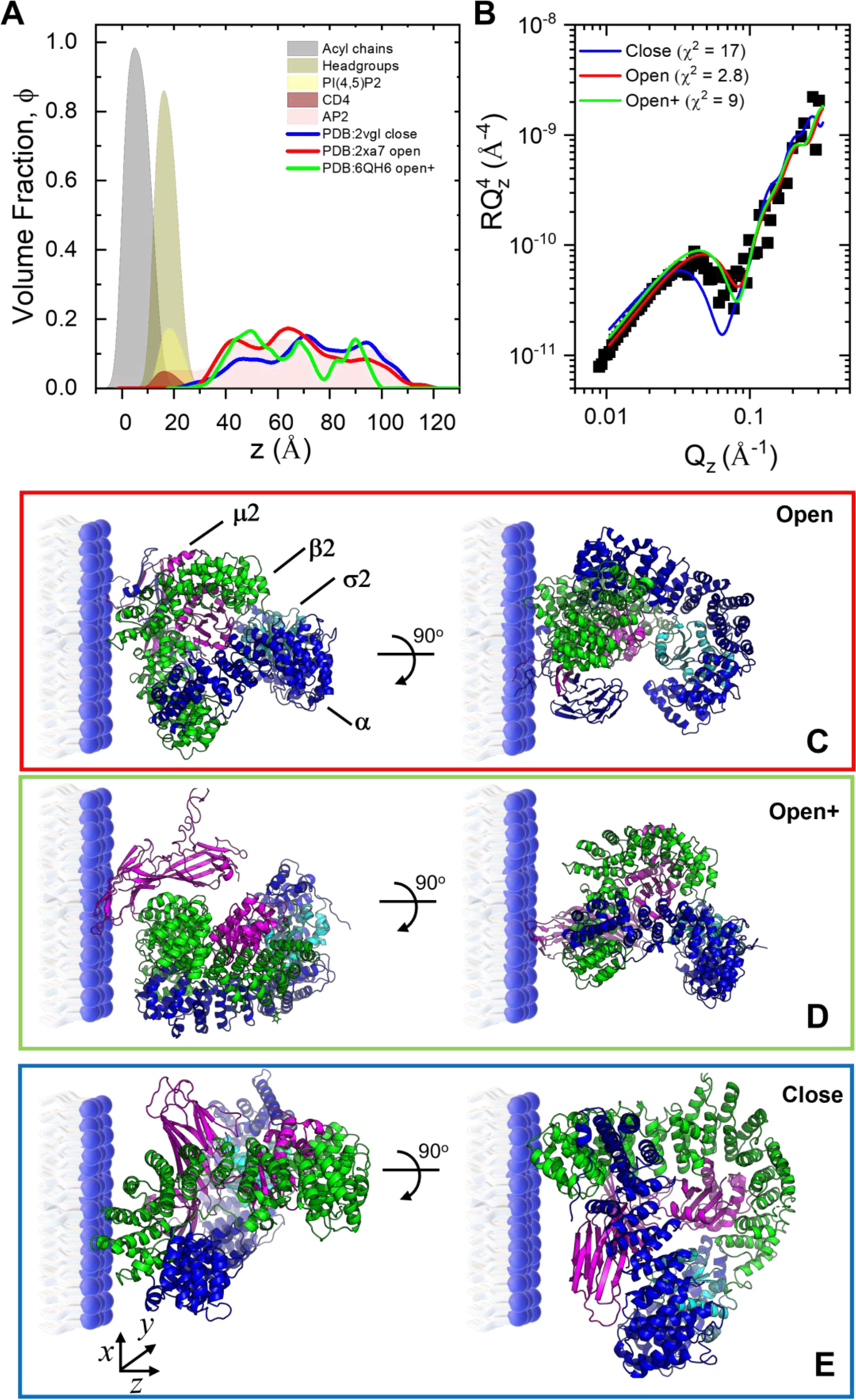
High resolution SNR data analysis using rigid-body modelling with AP2 crystal structures identifies open conformation as the preferred orientation and conformation of AP2 bound to cargo and PtdIns(4,5)P_2_ enriched monolayers. (A) Volume fraction profiles for open (PDB 2ax7) and close (PDB 2vgl) conformation of AP2. (B) Experimental SNR profiles in NRM compared to theoretical profiles calculated for open (red, C), open+ (green, D) and close (blue, E) conformation, respectively, based on the best possible orientations of the protein bound to the lipid monolayer. Heat maps evaluating the different protein orientations for open, open+ and close conformations as a function of the α and β Euler angles for intrinsic rigid body rotations of AP2 are shown in Figures S15-S17, respectively.

With regards to AP2 interacting with cargo-presenting PM model membranes, the calculated SLD profiles derived from open, open+ ^26^ and closed conformations were strikingly different, with the former always having better fits than the other two (**Figure 5 A, B** and **Figure S18**). In the most favourable scenario of an open AP2 conformation, the optimum position was found at a membrane head group distance of 10 ± 4 Å and 13 ± 4 Å, respectively for TGN38 and CD4 (**Figure 6C**). Consequently, AP2 exhibits minimal lipid interactions in comparison to the membrane lacking cargo (**Figure S19**. The protein’s orientation is consistent with the presentation of an open AP2’s PtdIns(4,5)P_2_ and cargo binding sites to the model PM (**Figure S18**). The surface coverage of AP2 for TGN38- and CD4-presenting PM was 51% and 24 % respectively, which agrees with tensiometry binding data (**Figure 6B**). Analysis of the cargo-less model membranes identified a similar open conformation of AP2, although the surface coverage is reduced to 12 % (**Figure 6A, B**). These observations agree with cryo electron tomography studies that identified an open conformation of AP2 on membranes enriched in PtdIns(4,5)P_2_ with or without TGN38 peptidolipids ^25,46^.

**Figure 6.**
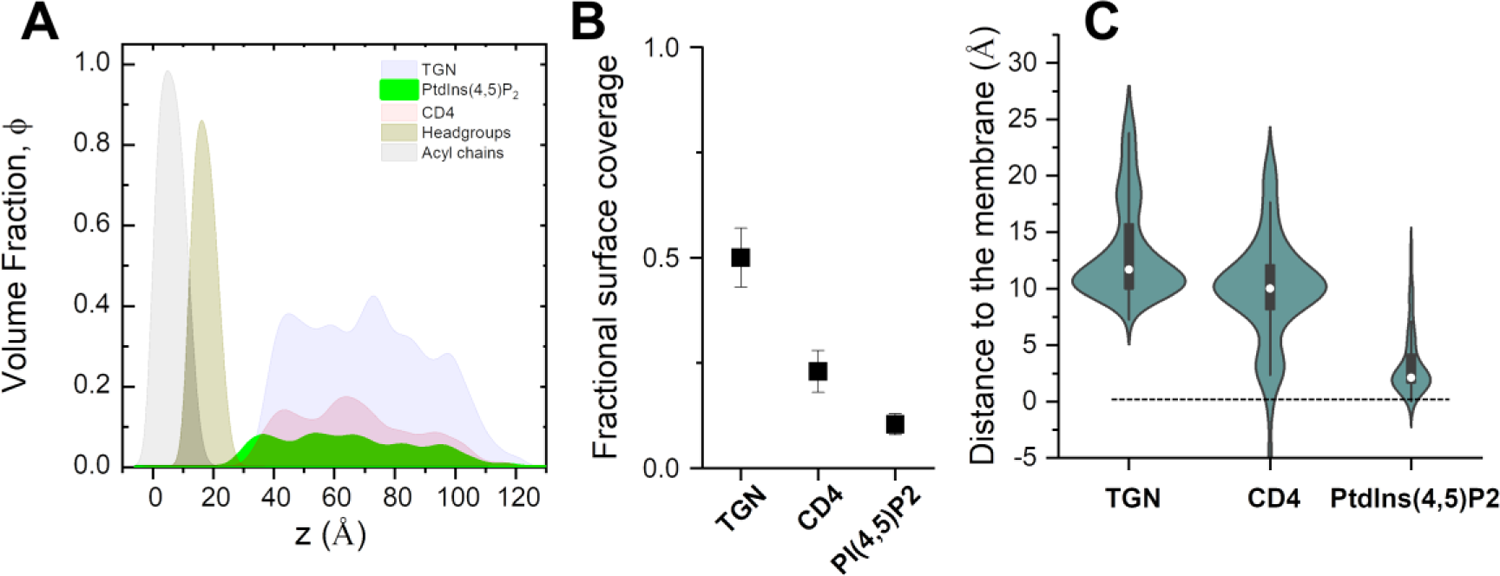
Analysis of high-resolution SNR data reveals that the orientation of open AP2 binding with PM models remains consistent regardless of cargo, with surface coverage being the sole variable. (A) Calculated volume fraction profiles corresponding to the preferred orientation of open conformation AP2 bound to PtdIns(4,5)P_2_-enriched monolayers in absence and presence of TGN38 or CD4 peptides. (B) Surface coverage and (C) Violin plots showing the distance to the interface were calculated by fitting the theoretical and experimental SLD values for each orientation of the protein in open conformation (PDB: 2xa7). The black vertical line indicates the interquartile range, the box plot, shows the ends of the first and third quartiles and the central dot the median value (n=256).

## Discussion and conclusion

Clathrin-mediated endocytosis (CME) is the primary mechanism through which eukaryotic cells selectively internalize proteins into the endosomal system. In solution, AP2 adopts a functionally closed (‘locked’) conformation, which precludes access to its cargo binding sites and simultaneously blocks all four of its PtdIns(4,5)P_2_-binding sites on a planar PM interaction ^46^. Consequently, on membrane binding, a large-scale conformational reorientation of its subunits is required for AP2 activation. Here, by using a battery of biophysical methods on bottom-up, minimal, *in vitro* membrane model systems, it was possible to address, under physiological conditions, the time-resolved recruitment and activation of AP2 by different PM models. Synergic interactions between adaptor protein and membrane lead to each molecularly adapting to each other.

With regards to the membrane, even though AP2 makes minimal lipid interactions, binding to the model PM monolayer induces a fluid-to-solid transition of the lipid leaflet, which is characterized by a larger viscosity and a finite shear elasticity. In contrast to cargo-less membrane, the increased binding of AP2 to cargo-presenting lipid monolayers leads to a drastic reduction in phospholipid mobility. Since AP2 binds specifically to PtdIns(4,5)P_2_, these rheometry data can be interpreted as AP2-induced clustering of PtdIns(4,5)P_2_, as previously observed with other PtdIns(4,5)P_2_ binding proteins ^47,48^. Below the percolation fraction ϕ_c_ = 0.73, where G’ = G’, the diffusion of AP2 and PtdIns(4,5)P_2_ is restricted to small, isolated liquid-phase domains (as observed in Figure 2). Above ϕ_c_ = 0.73, the membrane transitions from a fluid to a solid state, which occurs independently from clathrin binding. This result clarifies an intriguing previous study with rheometry that had focused on concurrent addition of clathrin and their associated adaptor proteins ^49^. The AP2 - PtdIns(4,5)P_2_ cluster network would tentatively be better able to resist shear deformation induced by clathrin polymerization, and subsequent CCV formation. Indeed, here we show for the first time that the formation of these 2D entropic networks might be therefore closely related to the regulation of plasma membrane mechanisms to incorporate and polymerize clathrin molecules to build endocytic pits.

With regards to AP2 adaptor, the cargo not only stabilized the open (“active”) conformation of AP2 ^11,24,40^, but also its presence on the plasma membrane. Intriguingly, our SNR structural data indicate that cargo containing either tyrosine Yxxφ motif and dileucine [DE]xxxL[LI] motifs may not play a critical role in AP2 “opening” conformational change, as interactions with PtdIns(4,5)P_2_-only PM is sufficient for full activation of AP2. This is in line with a recent tomography analysis, which had focused on carefully selected population of AP2 molecules binding to membranes, with and without TGN38^46^. However, in this current study, the inclusion of time-resolved kinetic data highlighted the AP2’s membrane surface dwell time, and resultant surface coverage, both significantly increase in the presence of cargo. We hypothesize the observed increases in AP2 residence time at the PM would allow sufficient time for clathrin to bind and therefore for productive nucleation of clathrin coated pits to proceed. A similar effect was observed *in vivo* by comparing wild type and mutant AP2 (the latter unable to bind specific cargo)^23^.

In conclusion, this *in vitro* analysis has yielded thermodynamic and structural insights into normal and aborted CME. It moreover provides proof-of-principle that other intracellular transport processes can be successfully studied using these experimental techniques and this conceptual framework developed in soft matter physics. This is particularly pertinent as the vast majority of these complex formation events occur deep within the cell and therefore are not compatible to characterization by current *in cell* techniques, like TIRF-SIM microscopy.

## Experimental section

### Lipids

1,2-dioleoil-sn-glycero-3-phosphoetanolamine (DOPE), 1,2-dioleoil-sn-glycero-3-phosphocoline (DOPC), 1,2-dioleoil-sn-glycero-3-phospho-L-serine (DOPS), Brain L-α-phosphatidylinositol-4,5-bisphosphate (PtdIns(4,5)P_2_) and cholesterol (Chol), were purchased as powder from Avanti Polar Lipids (purity >99%, Alabaster, AL, USA). Stock solutions (1 mg/mL) of the lipid mixtures were prepared in chloroform stabilized with ethanol (purity 99.8%; Sigma-Aldrich, St. Louis, MO, USA). See **Supplementary Note 1** for further details. D_2_O (99.9% of isotopical purity) was purchased from Sigma-Aldrich and used as received. HKT buffer was 25mM HEPES, 125mM Potassium Acetate and 1mM DL-Dithiothreitol (DTT), pH (7.20±0.05). HKM buffer was HKT buffer supplemented with 5mM Magnesium Acetate.

### Peptide-lipid conjugation

Peptides were synthesized with amino-terminal cysteines, and covalently conjugated to 1,2-dipalmitoyl-sn-glycero-3-phosphoethanolamine-N-[4-(p-maleimidophenyl)butyramide^11^. The Yxxϕ-containing peptide TGN38 was derived from the cytoplasmic tail of TGN38 (CKVTRRPKASDYQRL) while the di-leucine peptide CD4 was derived from the phosphorylated cytoplasmic tail of CD4 (CHRRRQAERMS*QIKRLLSEK) (where * denotes phosphorylation) (**Table S3** and **Supplementary Note 2**).

### Expression and purification of AP2

Recombinant AP2 were prepared in 250mM NaCl 2mM DTT 10mM Tris pH 8.7 according to previous protocols (Collins et al., 2002a). Briefly, after induction with 0.2mM IPTG followed by over-night expression at 22°C in BL21 plyS *E. coli* grown in 2TY media, cells were lysed using a cell disruptor (Constant Systems). AP2 were initially isolated with GST-beads and cleaved overnight with thrombin. The resultant protein was then isolated on NiNTA-beads, washed with buffer supplemented with 10mM imidazole, and eluted from the beads with the same buffer supplemented with 300mM imidazole. After size exclusion chromatography on a Superdex 200 column (GE Healthcare) in 250mM NaCl 10mM Tris pH 8.7 1mM DTT, the AP-2 complexes were concentrated to >15mg/ml with vivaspin concentrators, and flash frozen in 10% glycerol, until further use.

### Lipid monolayer and Langmuir trough experiments

A Langmuir trough (Kibron, Helsinki, Finland) was used to measure the surface pressure (Π) – area per molecule (A) isotherm of lipid monolayers as well as to determine the increase in pressure (ΔΠ) observed after the injection of the protein AP2 in the bulk phase. The variation of surface pressure was recorded using a Wilhelmy plate made of filter paper. Temperature was maintained at 21.5 ± 0.5°C. Further experimental details are given in the **Supplementary Note 3**.

### Specular neutron reflectometry (SNR)

**SNR** elucidates the structure and composition of interfacial layers in the direction perpendicular to the plane of the interface. Experiments were performed on the time-of-flight reflectometer FIGARO at the ILL. Two different angles of incidence (θ1 = 0.6° and θ2 = 3.7°) and a wavelength resolution of 7% dλ/λ, yielding a momentum transfer, Q_z_ = (4π/λ) sinθ, range from 0.007 to 0.25 Å^-1^ (the upper limit being limited by sample background) were used to perform the measurements and investigate the structure of the lipid monolayers upon AP2 interaction. Lipid monolayers were prepared in a Langmuir trough at a surface pressure of Π = 25 ± 1 mN/m. After SNR measurements of the lipid monolayer, AP2 was injected under the monolayer by a Hamilton syringe to a final bulk concentration of 10 nM. Two different isotopic solvent contrasts (100% D_2_O and 8.1% D_2_O (v/v %) respectively) were used for the characterization. Subsequent data analysis was performed using AuroreNR^50^ and Motofit^51^ software (see **Supplementary Note 8**). Optimization of model parameters was done by the combine use of a genetic algorithm for efficient search of the parameter space, and a Levenberg-Marquadt non-linear least-square algorithm for a final refinement of the fitting parameters. The quality of the fits was reported as reduced χ^2^.

### Imaging lipid monolayers by epifluorescence microscopy

Lipid monolayers were doped with 0.1 mol % BODIPY-TMR PtdIns(4,5)P_2_ and observed using an inverted bright-field microscope (Nikon Eclipse) with an extra-long working distance (WD 3.7–2.7 mm, NA 0.60) objective of 40× magnification. A CMOS camera (AVT Marlin F-131B) working at 30 frames per second was used to record a 160 μm × 120 μm field of view. Experimental limitations (drift motion, long-working distance and the diffraction limit of light) prevented an ideal visualization below the micron range.

### Imaging supported lipid monolayers by liquid AFM

Supported lipid monolayers were prepared by transferring monolayers from the Langmuir trough onto glass coverslips using the Langmuir−Schaeffer method. AFM images of fluid-phase lipid monolayers were taken using a Cypher S microscope with a bluedrive photothermally excited cantilever Olympus OMCL-AC240TS (240 x 40 μm, large x width) working at a resonance frequency of 70 kHz (in air, and, approximately 24 kHz in liquid) and an elastic constant of 1.2 N/m. The images were taken with a scan speed of 1Hz in tapping mode with 256 x 256 pixel resolution and 5 μm in size. Temperature and humidity conditions were stable. A droplet of HKM solution with an AP2 concentration was spread on the Langmuir-Schaeffer film of lipid, being visualized before droplet evaporation at a constant temperature.

### Interfacial Shear rheology

The mechanical response of the monolayers under shear deformation was measured by means of a homemade magnetic microwire interfacial shear rheometer driven by a mobile magnetic trap. The experiments were oscillatory with a frequency of 0.5 rad/s. The shear strain applied to the monolayer remained consistently below 1% throughout the duration of the experiment, where it is expected to be operated within the linear regime. The subphase and surface drags on the microwire were decoupled by numerically solving the flow field at the shear channel, and the surface dynamic moduli, G’ and G’’, were calculated from the surface velocity gradient at the contact line with the microwire ^52^. ^53^ (see **Supplementary Note 4,** for more details)

## Data Availability

The neutron data that supports the findings of this study are openly available in the Institute Laue-Langevin data portal: http://doi.ill.fr/10.5291/ILL-DATA.8-02-780, http://doi.ill.fr/10.5291/ILL-DATA.8-02-795 and http://doi.ill.fr/10.5291/ILL-DATA.8-02-804. The additional raw data supporting the results of our study area available within the article and its supplementary materials. The data analysis tools utilized are available upon request from the corresponding author, A.M.

## Acknowledgments

The authors thank the Institut Laue-Langevin for the allocation of beamtime and the Partnership for Soft Condensed Matter (PSCM) for the lab support. A.M. acknowledges the financial support from The Wellcome Trust Institutional Strategic Support Fund (ISSF) for a Junior Interdisciplinary Fellowship; from MICINN under grant PID2021-129054NA-I00, the Department of Education of the Basque Government under grant PIBA-2023-1-0054 and from the IKUR Strategy under the collaboration agreement between Ikerbasque Foundation and Materials Physics Center.

## Author Contributions

A.M., N.R.Z., D.O and P. C. conceived the study and planned the experiments. N.R.Z. expressed and purified all proteins. A.M., N.R.Z., J.F.G.M., P.G. and R.C. performed neutron experiments. A.M, J.F.G.M., P.S.P., J.T., A.S., J.C.T, D.P., carried out the rest of the experiments. AM processed all the experimental data and performed the analysis. A.M., N.R.Z., D.O. and P.C. wrote the manuscript. All authors provide critical feedback and helped shape the research, analysis and manuscript.

