## Supplementary Information for "Combined thermodynamic and time-resolved structural analysis of interactions between AP2 and biomimetic plasma membranes provides insights into clathrin-mediated endocytosis"

**Supplementary Notes:**

**Figure S10. Specular Neutron Reflectometry on Langmuir lipid monolayers**. **(A)** Neutron reflectivity data of the different hydrogenated lipid monolayers in $D_2O$ and ACMW buffers, respectively. The fitting curves of TGN lipids (blue), CD4 (red) and PIP2 (geen) are also shown. The figure is displayed on a $RQ^4$ scale to show the agreement between the experimental and theoretical data at higher q-values. (**B)** Scattering length density, SLD, profiles corresponding to fits are plotted in **A**. **C** Volume fraction profiles normal to the interface of monolayers to highlight the distribution of acyl chains and headgroups. ....................................................................................... 31

**Figure S11.** (**A**) Reflectivity profiles of DOPC:DOPE:PtdIns4,5$P_2$:CD4 (66.5:20:10:3.5) in 4 contrasts: $H_2O$ (dark blue circle), ACMW (blue triangles), SiMW (violet squares) and $D_2O$ (pink diamonds). The relative fits are presented as lines. (**B**) shows the SLD profiles perpendicular to the interface. (**C**) reports the corresponding volume fraction profiles perpendicular to the interface, showing the contribution of silicon oxide (orange line), aliphatic tails (black line), hydrophilic headgroups (magenta), and water (cyan line). The volume fraction related to CD4 peptide moiety is shown as light pink area.. 32
**Figure S15 Atomic structures of AP2 interacting with CD4 enriched monolayers**. Energy minimized crystal structures from PDB 2ax7 and from 2vgl. used to position and orient AP2 component of the experimentally-derived SLD profile. The orientations were parameterized using the Euler angles $\alpha$ and $\beta$ (following an x-y-z extrinsic rotation scheme). An ensemble of 246 different orientation trialled, with 5$^{o}$ increments in $\alpha$ and $\beta$. A and B show colormaps of the preferred orientations for 2xa7 and 2vgl, whilst c and d the respective SLD profiles generated for AP2 as a function of the distance to the interface. ........ 35
**Figure S16. Atomic structures of AP2 interacting with TGN enriched monolayers**. Energy minimized crystal structures from PDB 2ax7 and from 2vgl. used to position and orient AP2 component of the experimentally-derived SLD profile. The orientations were parameterized using the Euler angles $\alpha$ and $\beta$ (following an x-y-z extrinsic rotation scheme). An ensemble of 246 different orientation trialled, with 5$^{o}$ increments in $\alpha$ and $\beta$. A and B show colormaps of the preferred orientations for 2xa7 and 2vgl, whilst c and d the respective SLD profiles generated for AP2 as a function of the distance to the interface. ........ 36
**Figure S17**. **Atomic structures of AP2 interacting with PtdIns(4,5)P2 enriched monolayers**. Energy minimized crystal structures from PDB 2ax7 and from 2vgl. used to position and orient AP2 component of the experimentally-derived SLD profile. The orientations were parameterized using the Euler angles $\alpha$ and $\beta$ (following an x-y-z extrinsic rotation scheme). An ensemble of 246 different orientation trialled, with 5$^{o}$ increments in $\alpha$ and $\beta$. A and B show colormaps of the preferred orientations for 2xa7 and 2vgl, whilst c and d the respective SLD profiles generated for AP2 as a function of the distance to the interface. ........ 37
**Figure S18.** Comparison of the favored orientations of AP2 in presence of CD4 (A), TGN (B) and PtdIns(4,5)P2 (C) enriched monolayers for open (2xa7) and close (2vgl) conformations with the experimental scattering length densities (SLDs) obtained by NR. ........ 38
**Figure S19**. (A) Comparison of *CD4, TGN and PtdIns(4,5)P2 enriched monolayers for open (2xa7) conformation with the experimental SLDs. Corresponding values of volume fractions (B) and anchor distance (c) obtained from the fitting using the theoretical SLD profiles obtained.* ........ 38
**Figure S20. SNR – low $Q_z$ analysis**. (A) Reflectivity profiles in the region 0.01 Å$^{-1}$ < $Q_z$ < 0.03 Å$^{-1}$. Color bar indicates the evolution of time after AP2 is injected in the

subphase. (B) Selected reflectivity profiles fitted with a one-layer model (straight lines). (C) corresponding SLD profiles as a function of the distance to the interface. (D) AP2 recruited vs time obtained from the fitting of the experimental profiles shown in (A) based on the SLD distribution plotted in C. ................................................................ 39
**Figure S 21. SNR and QCM-D data of Lipid bilayers enriched in PtdIns(4,5)P2 and CD4 with AP2**: (A) QCM-D data: Frequency shift and dissipation shift plots (corresponding to the 3rd overtone, n=3) showing lipid vesicles (DOPC, DOPE, PtdIns(4,5)P2 and CD4) adsorption and fusion kinetics: lipid vesicles and subsequent AP2 injection. (B)Volume fraction profiles of solid supported lipid bilayers derived from SNR and SLD profiles plotted in Figure S11. (C) Scheme of the resulting AP2 low-resolution structure bound to lipid bilayers. ................................................................ 41

### 1. Extended Information about samples

***Table S1*** *Lipid nomenclature, molecular formula and weight.*

| **Name** | **Abbrev.** | **Molecular Formula** | **Molecular weight (g/mol)** |
|---|---|---|---|
| **1,2-dioleoil-sn-glycero-3-phosphoetanolamine** | DOPE | C41H78NO8P | 744.034 |
| **1,2-dioleoil-sn-glycero-3-phosphocoline** | DOPC | C44H84NO8P | 786.113 |
| **1,2-dioleoil-sn-glycero-3-phospho-L-serine** | DOPS | C42H77NO10P | 810.025 |
| **L-α-phosphatidylinositol-4,5-bisphosphate** | PtdIns(4,5)$P_2$ | C47H94N3O19P3 | 1096.385 |
| **Cholesterol** | Chol | C27H46O | 388.67 |

***Table S2*** *Lipid composition of Langmuir used in this study.*

| Molar % | DOPE | DOPC | DOPS | PtdIns(4,5)$P_2$ | Chol | CD4 | TGN38 |
|---|---|---|---|---|---|---|---|
| **PtdIns(4,5)$P_2$** | 32.4 | 30.7 | 9.9 | 14.7 | 12.4 | - | - |
| **CD4-PtdIns(4,5)$P_2$** | 31.5 | 29.8 | 9.7 | 14.3 | 12.1 | 2.6 | - |
| **TGN38 - PtdIns(4,5)$P_2$** | 31.3 | 29.6 | 9.6 | 14.1 | 12.0 | - | 2.5 |

*Table S3* *Sequence of amino acids of the peptides studied.*

| Sample | Sequence | Mw (kDal) | MW (Lipid+peptide) (kDa) |
|---|---|---|---|
| **CD4** | CHRRRQAERMSQIKRLLSEK | 2.562 | 2.776 |
| **TGN38** | CKVTRRPKASDYQRL | 1.837 | 3.481 |

***Table S4*** *Different components of AP2 with the respective structure PDB ID*

| Protein | PDB Entry | Molecular Weight, kDa |
|---|---|---|
| AP2-Core : Open | 2XA7 | 205 |
| AP2-Core : Close | 2VGL | 203 |
| β2-Adaptin Appendage | 1E42 | 29,06 |
| α-Adaptin Appendage | 1B9K | 26,99 |
| AP2 full length * | x | 287 |

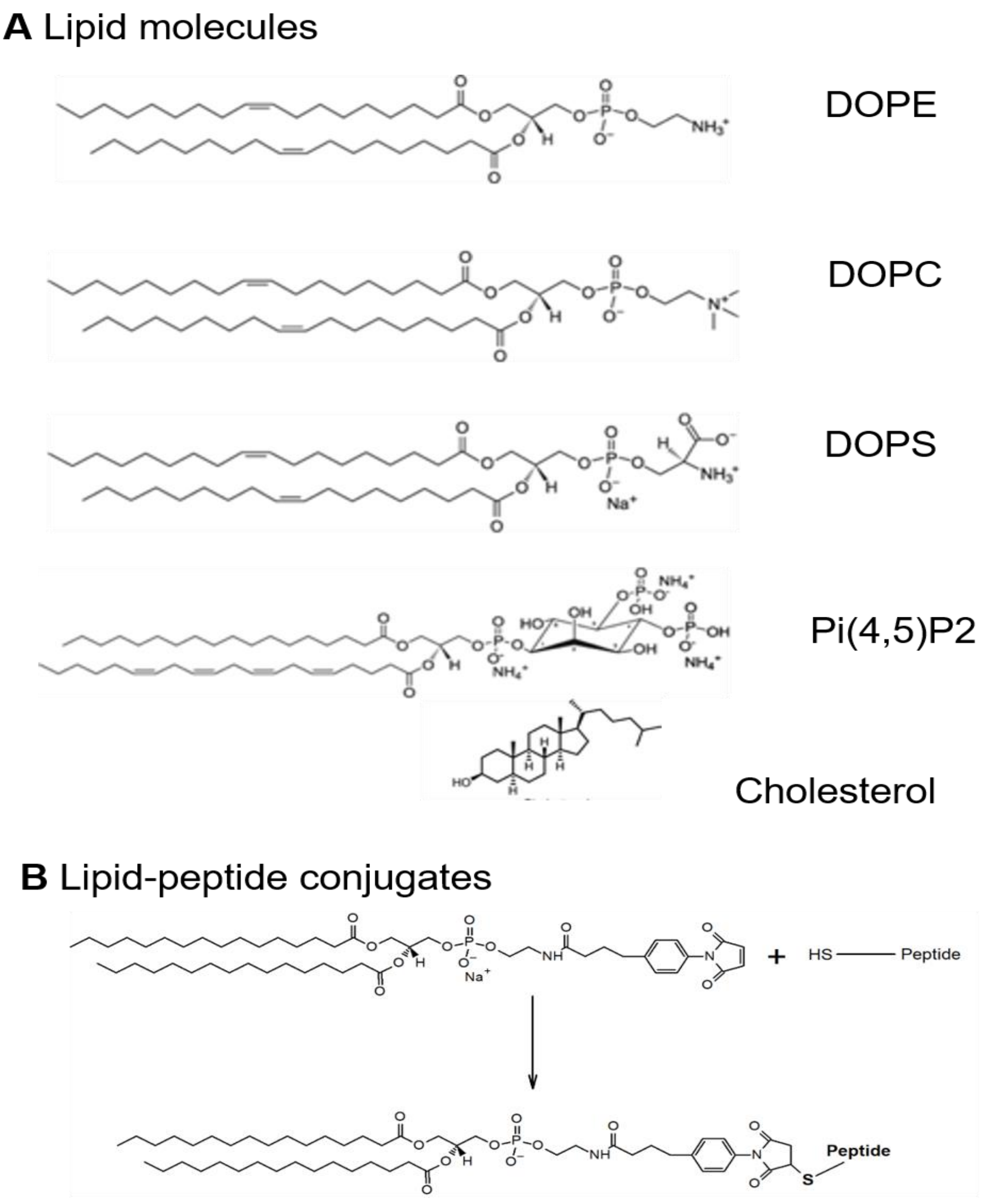

**Figure S1** (A) Lipid composition. (B) Schematic representation of the synthesis route in the conjugation of a peptide sequence with a synthetic lipid. The conjugation is achieved through the reaction of the cysteine, present in the N-terminus, with the maleimide group of 16:0 MPB PE lipid.

### 2. Cargo molecules synthesis

Peptides sequences, containing the sorting recognition motifs from the cytoplasmic tails of TGN38 and phosphorylated CD4, were synthesized with a cysteine moiety in the amino-terminal. The quality of the produced peptides was evaluated with mass spectrometry and reverse-phase chromatography. After reaching the quality criteria desired for the peptides, a conjugation reaction was performed where the peptides were covalently conjugated to a synthetic lipid, 1,2-Dipalmitoyl-sn-Glycero-3-Phosphoethanolamine-N-[4-(p-maleimidophenyl)butyramide] commonly known as MPB PE, as shown in **Figure S1**. The conjugation is achieved through the reaction of the cysteine moiety, present in the N-terminus, with the maleimide group of 16:0 MPB PE lipid. The final solutions of the lipopeptide were collected after extraction, dried under nitrogen, resuspended in chloroform:methanol (2:1) and stored at -20ºC.

The amino acid sequence and respective molecular weight of each peptide, before and after conjugation with 16:0 MPB PE, is reported in **Table S3**.

### 3. Langmuir trough experiments

#### Lipid monolayer fabrication

A Langmuir trough with a maximum area of 166.4 $cm^2$ equipped with two dependent barriers (Kibron, Helsinki, Finland) was used to measure the surface pressure (Π) – area (A) isotherms of lipid monolayers (See **Figure S2**). Π is recorded using a Wilhelmy plate made of chromatographic paper (Whatman CHR1). In detail, a reduction in the interfacial tension of the bare air/buffer interface $\gamma_0$ (72.5 ± 0.5 mN/m at 21.5ºC) by the presence of a lipid monolayer with an interfacial tension $\gamma$ is defined as a surface pressure $\Pi = \gamma_0 - \gamma$, in analogy to the bulk osmotic pressure.

After careful cleaning, the trough was filled with HKM buffer (25mM HEPES, 5mM Magnesium Acetate ($Mg(CH_3COO)_2$) 4H2O, 125mM Potassium Acetate ($CH_3CO_2K$) and 1mM DL-Dithiothreitol (DTT)), and a 0.1 mg/mL lipid solution in chloroform was spread over the subphase using a Hamilton micro-syringe with a precision of ±1 µL. After the chloroform was evaporated for about 20 min, the variation of surface pressure during compression was recorded at a barrier speed of 5 $cm^2$/min. The subphase temperature was maintained at 21.5 ± 0.5 °C by making thermostatic water flow through jackets at the bottom of the trough. After the evaporation of the chloroform and the stabilization of the monolayer, peptides were injected underneath the lipid monolayer in the bulk phase.

### Surface pressure – area isotherms and lateral compressibility of the monolayer.

The evolution of lateral pressure as a function of the average area available for each lipid molecule provides insightful information about the lipid phase behavior. Indeed, it allows to distinguish if a lipid monolayer is in a liquid expanded (LE, fluid-like) phase or in a liquid condensed (LC, gel-like) phase. Furthermore, from the slope of the $\Pi$-A isotherm it is possible to determine the lateral compressibility of the monolayer, i.e., its mechanical resistance against a dilational deformation in terms of the compressional elastic modulus of the film.

### Binding affinity determination

The binding constant of AP2 with the lipid monolayer was calculated by measuring the increase in pressure, $\Delta\Pi = \Pi_\infty - \Pi_0$, observed after the injection of the peptides in the bulk phase at an initial surface pressure $\Pi_0 = 25 \pm 1$ $mN{\cdot}m^{-1}$, to mimic the inner leaflet of the plasma membrane, until the surface pressure reached a plateau, $\Pi_\infty$ as shown **Figure S3.** The differences in the surface pressure $\Delta\Pi$, which depend on the amount of AP2 injected, were normalized with respect to the maximum value of difference obtained ($\Delta\Pi_\infty$). The obtained $\Delta\Pi/\Delta\Pi_\infty$ versus peptide concentration profiles were analyzed with a Langmuir adsorption model based on an adsorption equilibrium equation and mass balance between the monolayer and the FPs, assuming that the degree of normalized surface pressure increase is proportional only to the coverage fraction of peptides interacting with the monolayer φ:

$$\varphi = \frac{\Delta\Pi}{\Delta\Pi_{max}} = \frac{C^n}{C^n + K_D} \quad \textbf{(S1)}$$

where C is the bulk concentration of the peptide, n is the Hill coefficient and $K_D$ is a dissociation constant.

### Kinetic Binding analysis of AP2 to TGN model PM.

From the Stokes-Einstein equation, the diffusion coefficient of AP2 can be calculated as D = kBT=6πη*a*, assuming a spherical shape with a radius *a* = 50Å, based on AP2 crystal structures. The value obtained (D = $4.3 \cdot 10^{-11}$ $m^2/s$) can be comparable with the one from the empirical Polson equation D = $\beta Mw^{-1/3}$= $4.7 \cdot 10^{-11}$ $m^2/s$, with Mw = 200 kDa and β = $2.74 \cdot 10^{-9}$ (Polson, 1967). Before analyzing dynamic data by using a mechanism based on a kinetic barrier, we confirmed that mass transport by diffusion was not limiting the adsorption of AP2. The characteristic time $t_d$ of a diffusion-controlled adsorption process was estimated from the following (Middelberg et al., 2000)

$$t_d = \frac{1}{D}\left(\frac{\Gamma_\infty}{C_{AP2}}\right)^2$$

where D is the AP2 diffusion coefficient in bulk, estimated by SE (and Polson equation), $C_{AP2}$ is the bulk AP2 concentration, and $\Gamma_\infty$ is the maximum interfacial coverage (calculated by neutron reflectometry). A characteristic diffusion time $t_d$ = 160 s is calculated based on the following values: $C_{AP2}$ = 45 nM, D = $4.7 \cdot 10^{-11}$ $m^2/s$, and $\Gamma_\infty$= 3.96 nMol/$m^2$. In view of this value of $t_d$, we can conclude the adsorption of AP2 in the interval of concentrations studied here is considerable slower than would be expected under diffusion control. We, therefore, consider the binding of AP2 to lipid monolayers enriched in TGN is not limited by diffusion, and we propose a mechanism based on a kinetic limitation. Similar values of $t_d$ were found for monolayers enriched in CD4.

Modelling the binding process based on a first order irreversible reaction, with no desorption of AP2 once linked to the lipid monolayer, the surface concentration of AP2, denoted as $\Gamma$ is, therefore, solely determined by the transition of the molecules over a kinetic barrier. The rate of increase in $\Gamma$ can be expressed as $d\Gamma = k\Gamma$, where *k* represents the rate constant of binding. Assuming a linear equation of state (*i.e.*, a linear relation between the interfacial binding of AP2 and the surface pressure $\Pi(t) = k_\theta\ \Gamma(t)$, where $k_\theta$ is a proportionality constant. Solving this Equation, normalizing it by $\Pi_\infty$, the steady state value of surface pressure at $t \rightarrow \infty$ and accounting for the non-exponential character of some experimental curves by including a stretched exponential $\beta$ as a representation of rate distributions we obtain **Equation 2** in the main text.

**Table S5**. Results of the kinetic binding analysis performed on TGN enriched monolayers.

| $C_{AP2}$ ± 5 / nM | $\beta$ ± 0.1 | t ± 0.1 / min | $k_d$ ± 0.005 / min$^{-1}$ |
|---|---|---|---|
| 20 | 1.0 | 63.7 | 0.016 |
| 37 | 1.0 | 50.7 | 0.020 |
| 45 | 0.9 | 38.9 | 0.026 |
| 50 | 0.8 | 26.3 | 0.038 |
| 100 | 0.6 | 12.6 | 0.079 |
| 170 | 0.5 | 4.5 | 0.140 |
| 500 | 0.4 | 2.2 | 0.448 |

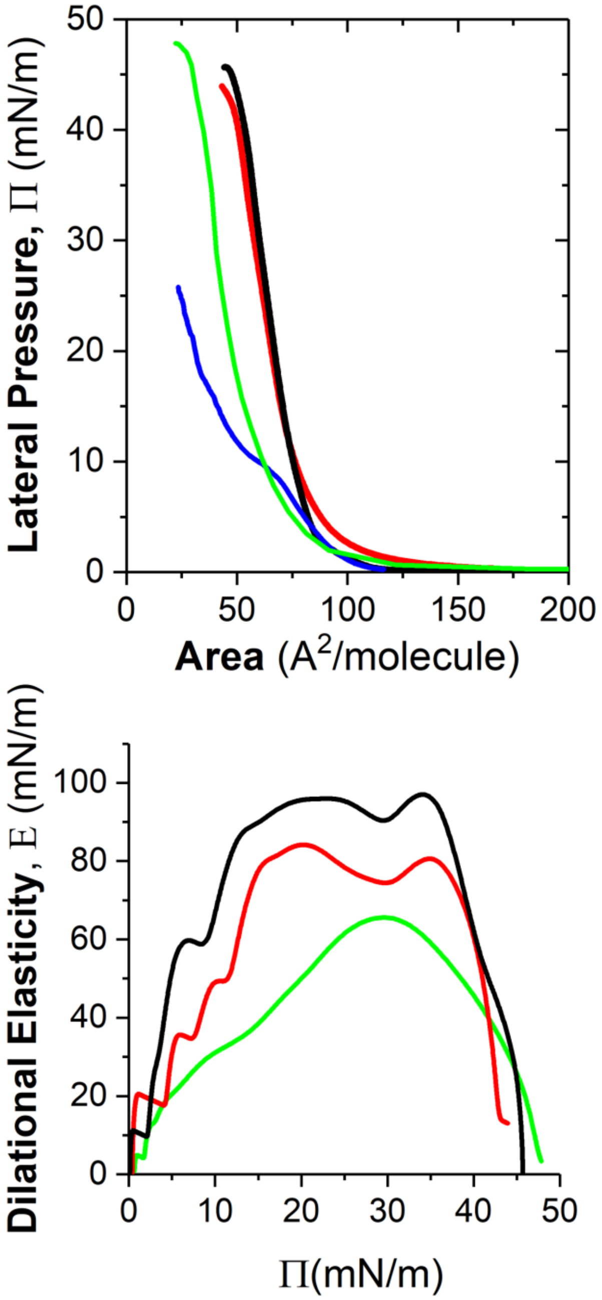

**Figure S2** (Top) Surface pressure – Area per molecule isotherms at T = 21.5 ºC of TGN mixture (black line), CD4 mixture (red), PIP2 mixture (green) and DOPE:DOPC:DOPS:Chol (blue) monolayers. (Bottom) Corresponding compressional elastic modulus obtained from the slope of the isotherm.

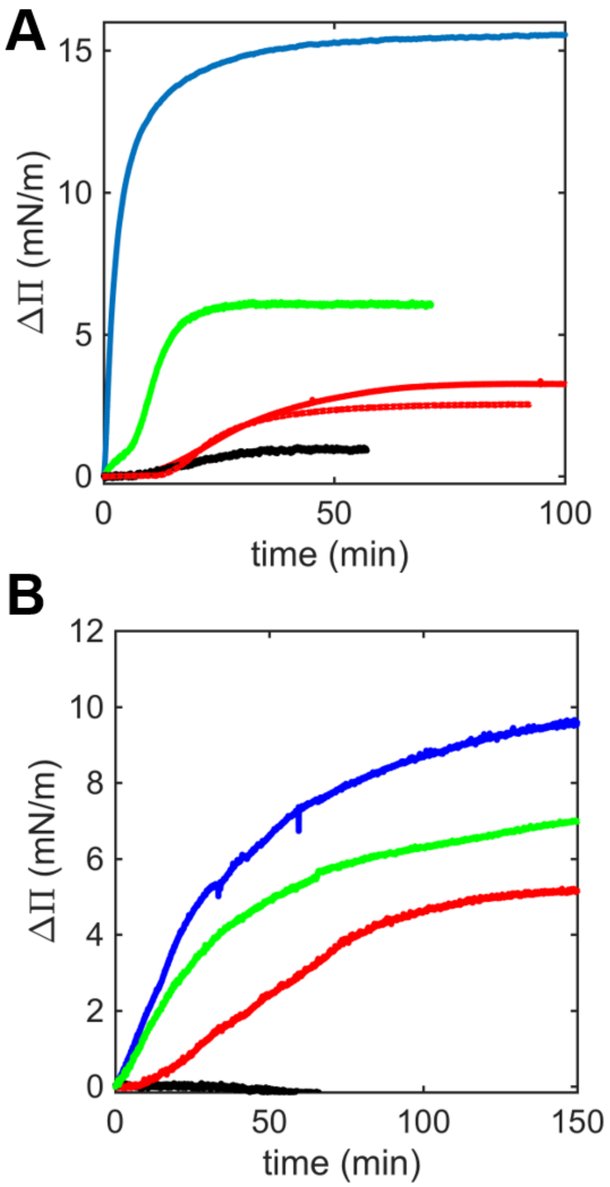

***Figure S3*** Experimental change in surface pressure $\Delta\Pi_\infty$ upon AP2 interacting with different lipid monolayers: (A) Comparison between CD4 (green line), TGN (blue) and PIP2 (red) enriched monolayers at Π=25mN/m. Control is a mixture of DOPE,DOPS,DOPC and Chol (black). (B) Comparison at increasing concentration of PIP2 up to 20 % mol.

**4. Interfacial Shear rheology**

The sample was placed in a custom-made PTFE container with 5 mm in depth, having a L*W shear channel and a S*S reservoir for sample preparation and surface pressure measurement purposes. A thin magnetic microwire (24.6 µm in diameter and ∼1 cm in length) was placed at the interface and used as a probe. The magnetic trap is placed above the shear channel and comprises two small permanent magnets that align the microwire along the channel's direction. The dynamics of the trap-microwire system is well represented by an elastic restoring force proportional to the relative displacement between the center of the microwire and the midpoint between the magnets. The elastic constant (trap strength) can be tuned by vertically displacing the magnets, where the elastic constant increases when decreasing the vertical displacement between the magnets and the microwire. Therefore, a good compromise between sensitivity and strength can be found in a wide range of surface dynamic moduli, allowing for the accurate measurement of G' and G'' values ranging from $\sim 10^{-8}$ N/m to $\sim 10^{-2}$ N/m.

The schematics of the Magnetic Needle Interfacial Shear Rheometer is represented in the Figure S4 (Top). The magnetic trap consists of two small permanent magnets attached on a platform controlled by two different linear stages, one driven by a servo motor that moves the platform vertically (y axis), and another one driven by a stepper motor that moves the platform horizontally (z axis). The magnetic field near the magnets makes a magnetic needle located at the interface (below the trap) experience, first, a centering torque that aligns the needle along the shear channel, and second, an elastic-like restoring force proportional to the relative displacement between the center of the needle and the midpoint between the magnets. When the magnetic trap performs an oscillatory motion with frequency $\omega$ along the z axis, the motion of the needle rapidly reaches the stationary solution given by an oscillatory motion with the same frequency $\omega$. Let us choose the phase reference such that the displacement of the probe and the trap are, respectively

$$z_p(t) = \mathrm{Re}\{z_{p,0}\, e^{i\,(\omega\, t+\delta)}\} = \mathrm{Re}\{z^*_{p,0}\, e^{i\,\omega\, t}\},$$

$$z_t(t) = \mathrm{Re}\{z_{t,0}\, e^{i\,\omega\, t}\},$$

where $z_{p,0}$ and $z_{t,0}$ are the probe's and trap's motion amplitudes, respectively, and $\delta$ is the phase lag between both oscillatory displacements. We can relate these two oscillatory displacements through the complex amplitude ratio

$$\mathrm{AR}^* = \frac{z_{p,0}}{z_{t,0}}\, e^{i\,\delta} = \frac{z^*_{p,0}}{z_{t,0}}.$$

The force balance equation for the probe reads

$$F_m(t) + F_{sub}(t) + F_{surf}(t) = \mathrm{m}\frac{d^2 z_p(t)}{dt^2},$$

where $F_m(t) = -k\left(z_p(t) - z_t(t)\right)$ is the restoring magnetic force, $k$ being the elastic constant of the magnetic trap, $F_{sub}(t)$ is the viscous drag on the needle from the liquid subphase, $F_{surf}(t)$ is the drag on the needle from the interface, and $m$ is the probe's mass. Combining the last three equations, we can relate the raw data acquired by the instrument, $AR^*$, with $F_{sub}(t)$ and $F_{surf}(t)$ as

$$\frac{1}{\mathrm{AR}^*} = 1 + \frac{1}{k}\left(\frac{F_{sub} + F_{surf}}{z_{p,0}^*} - m\omega^2\right).$$

The subphase and surface drags are coupled in a non-trivial way, but the problem can be deconvoluted by explicitly solving the flow field in the shear channel. We follow the numerical approach described in (Reynaert et al., 2008) to solve the Navier-Stokes equations accounting for the Boussinesq-Scriven condition at the interface, allowing us to tackle eventual non-linear velocity profiles at the interface. Hence, the separation of the subphase and interface contributions to the probe's dynamics is feasible: they can be calculated from the velocity gradients evaluated at i) the probe's surface in contact with the subphase and ii) the contact line between the probe and the interface, respectively. Since the flow at both the subphase and the interface are coupled, an iterative scheme [ref] is necessary to find the proper values of G' and G'' that fit the force balance equation of the probe to the experimentally measured values of $AR^*$. Such iterative scheme was performed with dedicated MATLAB routines and is thoroughly described and analyzed in (Sánchez-Puga et al., 2021).

In the experiments here reported, $\omega =$ rad/s and the acquired data for $z_p(t)$ and $z_t(t)$ were analyzed in the frequency domain through a DFT routine to obtain $AR^*$. If needed, the elastic constant of the magnetic trap, $k$, was increased to keep the amplitude of the probe's displacement fairly constant, yielding a strain amplitude consistently below 1% throughout the duration of the experiments, where it is expected to be operated within the linear regime.

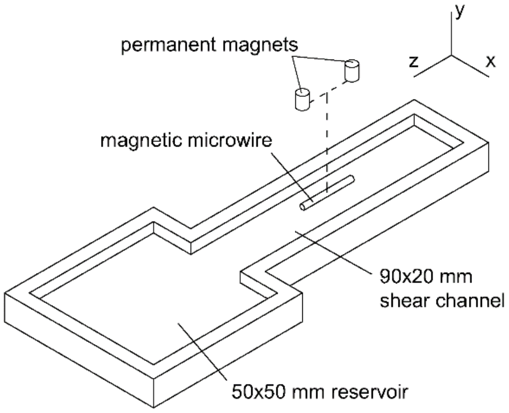

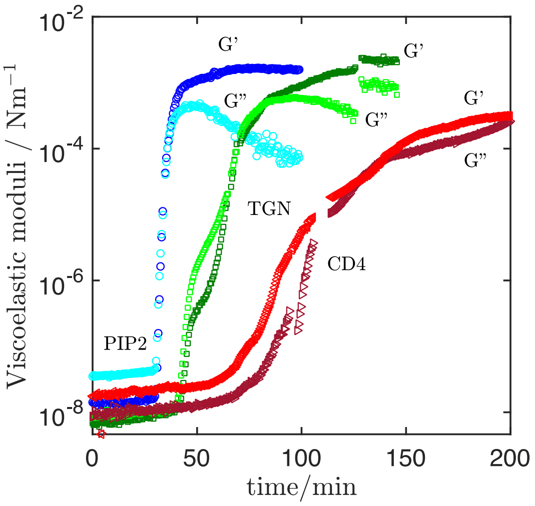

**Figure S4.** (Top) Schematics of the Magnetic Needle Interfacial Shear Rheometer. (Bottom) Experimental change in Shear surface moduli (G' and G'') upon AP2 interacting with different lipid monolayers: CD4, TGN and PtdIns(4,5)$P_2$ enriched monolayers at an initial surface pressure, Π=25mN/m.

### 5. Kinetic of binding follow by ellipsometry

Ellipsometry is a non-destructive optical technique based on the determination of the polarization changes that light undergoes when it is reflected at an interface. These changes are defined by the ratio of the overall Fresnel reflectivity coefficients of the parallel ($r_P$) and the perpendicular ($r_S$) components of the electric field, which are related to the ellipsometric angles Δ and Ψ. This relationship, known as ellipticity ρ, is defined by

$$\rho = {}^{r_p}/_{r_s} = tan\,\Psi e^{i\Delta}$$

The experiments were performed on a Picometer Light ellipsometer (Beaglehole Instruments, Kelburn, New Zealand) fitted with a He-Ne laser with $\lambda$ = 632 nm. The Langmuir trough (KIBRON, Helsinki, Finland) was coupled with the ellipsometer to measure the surface pressure of the lipid monolayer during the measurements of the ellipsometric angles. Lipid monolayers were deposited at the air/water (HKM buffer) interface. When an initial surface pressure of 25 ± 1 mN/m was achieved by compression with the barriers, AP2 was injected underneath the monolayer in the bulk phase and the ellipsometric angles $\Delta$ and $\Psi$ were measured, at an angle of incidence of 51$^{o}$ as a function of time (**Figure S5**). The increment observed in both $\Delta$ and $\Psi$ angles is proportional to an increase in either the refractive index and the thickness of the interfacial layer rationalized in either case by the growth in AP2 protein bound to the lipid monolayer.

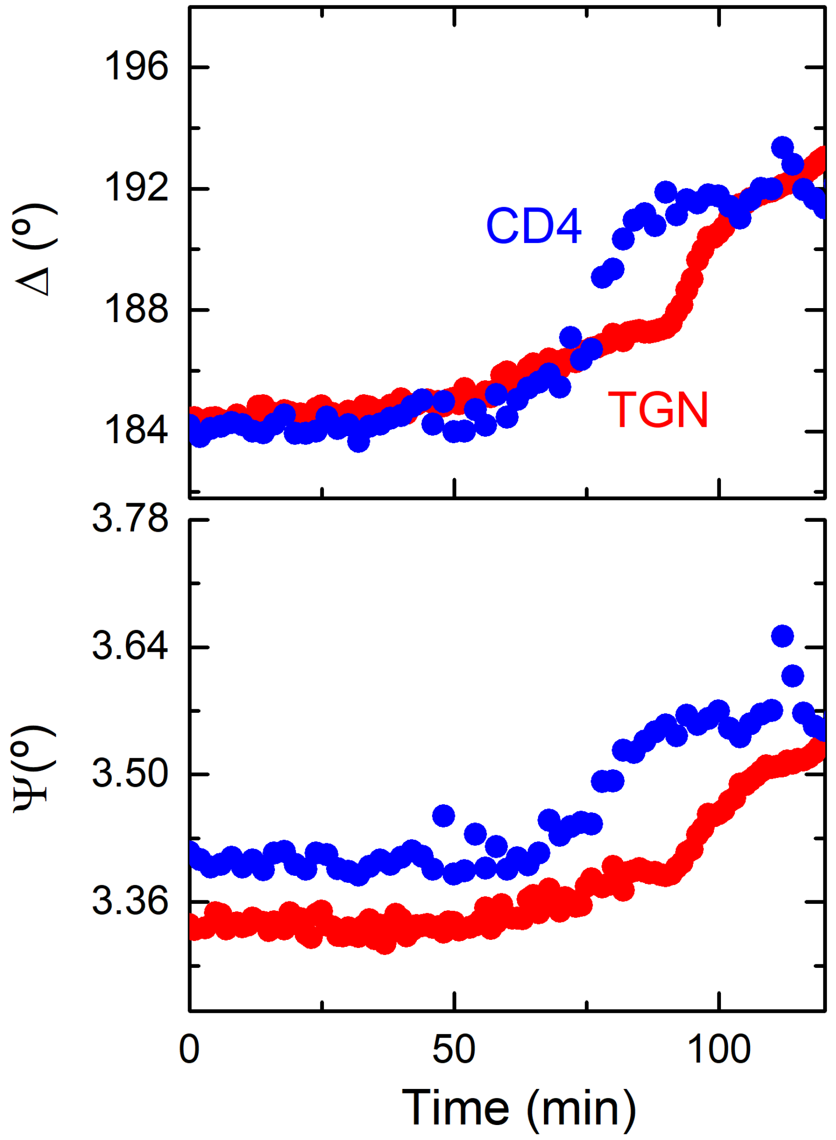

**Figure S5. Kinetic analysis of AP2 binding to a CD4 and TGN monolayers by ellipsometry.** Variation of ellipsometric angles Δ and Ψ with time after AP2 injected into the bulk phase. CD4 (blue) and TGN (red) enriched monolayers were formed at a surface pressure of 25 ± 1 mN/m. Both Δ and Ψ angles are proportional to the amount of AP2 protein bound to the lipid monolayer.

### 6. Epifluorescence microscopy on Langmuir monolayers

Lipid monolayers were doped with 0.1 mol % BODIPY-TMR PIP2 and observed using an inverted bright-field microscope (Nikon Eclipse) with an extra-long working distance (WD 3.7–2.7 mm, NA 0.60) objective of 40× magnification. A CMOS camera (AVT Marlin F-131B) working at 30 frames per second was used to record a 160 μm × 120 μm field of view. Experimental limitations (drift motion, long-working distance and the diffraction limit of light) prevented an ideal visualization below the micron range.

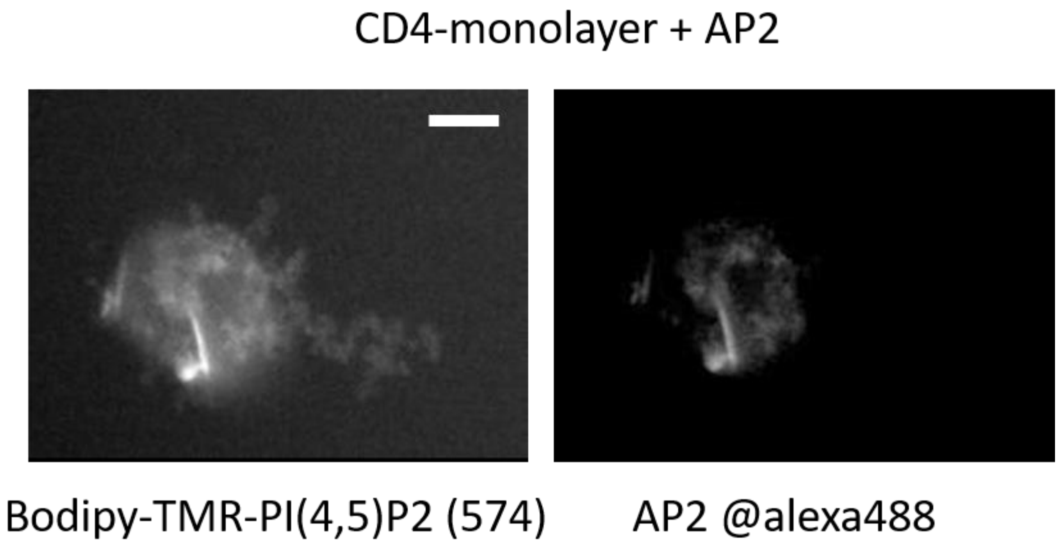

**Figure S6**. Simultaneous colocalization of AP2 and PtdIns(4,5)P2 clusters in Langmuir monolayers at the air water interface were reported by fluorescence images of CD4-PIP2 monolayer at the air/buffer interface labeled with 1 mol % Bodipy-TMR-PtdIns(4,5)P2 (left panel) interacting with AP2-labelled with Alexa Fluor 488 (right panel). Scale bar is 20 μm.

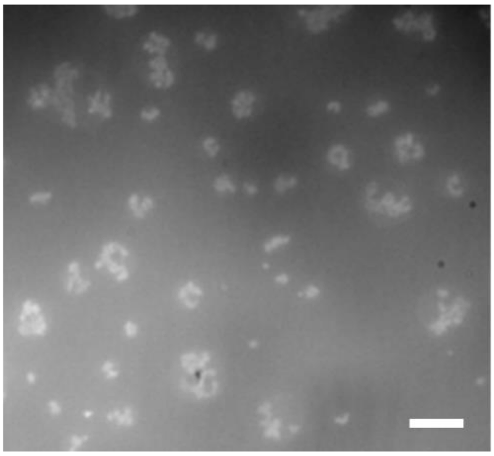

**Figure S7** Control experiment. PtdIns(4,5)P2 clusters in Langmuir monolayers at the air water interface were reported by fluorescence images of PtdIns(4,5)P2 monolayer at the air/buffer interface labeled with 1 mol % Bodipy-TMR-PtdIns(4,5)P2. Scale bar is 20 μm.

### 7. Atomic Force Microscopy (AFM) experiments on Langmuir-Schaeffer films.

Supported lipid monolayers were prepared by transferring monolayers from the Langmuir trough onto glass coverslips using the Langmuir−Schaeffer method. AFM images of fluid-phase lipid monolayers were taken using a Cypher S microscope with a blue-drive photothermally excited cantilever Olympus OMCL-AC240TS (240 x 40 μm, large x width) working at a resonance frequency of 70 kHz (in air, and, approximately 24 kHz in liquid) and an elastic constant of 1.2 N/m. The images were taken with a scan speed of 1Hz in tapping mode with 256 x 256 pixel resolution and 5 μm in size. Temperature and humidity conditions were stable. A droplet of HKM solution with an AP2 concentration was spread on the Langmuir-Schaeffer film of lipid, being visualized before droplet evaporation at a constant Temperature. In this section, supporting AFM images of Langmuir-Schaeffer layers are included.

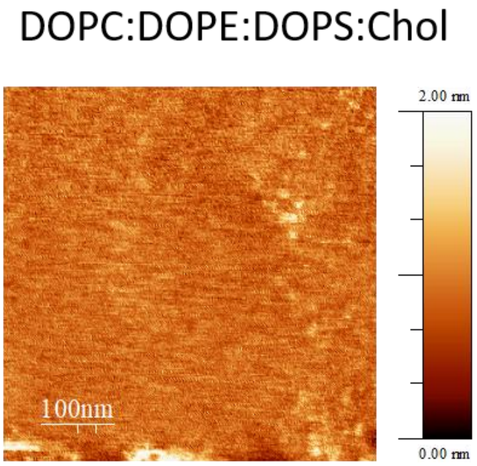

**Figure S8** Transferred lipid DOPC:DOPE:DOPS:Chol monolayers immersed in HKM buffer were examined by fluid-phase, tapping-mode AFM.

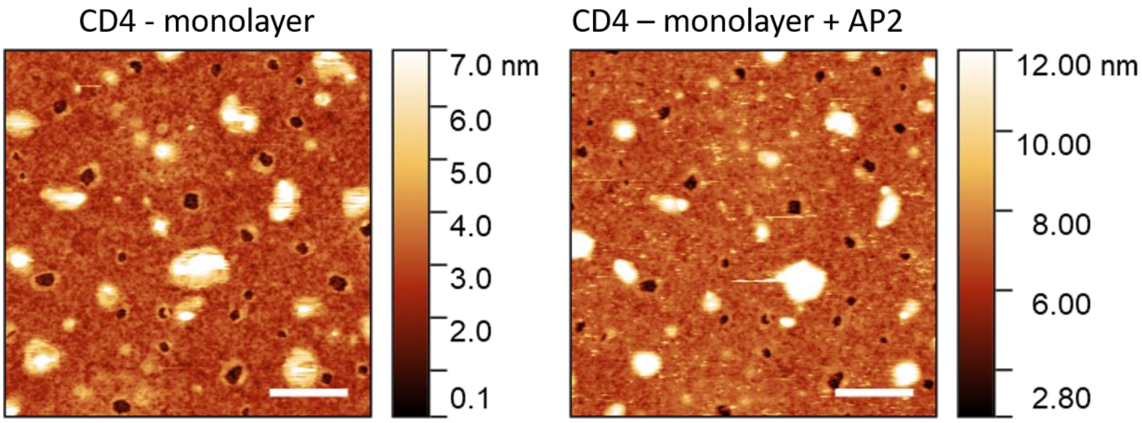

**Figure S9** Transferred lipid monolayers enriched in CD4 before and after incubation with AP2 examined by fluid-phase, tapping-mode AFM. Scale bars are 100 nm.

### 8. Specular Neutron reflectivity (SNR)

#### Definition

SNR elucidates the structure and composition of interfacial layers in the direction perpendicular to the plane of the interface. A 1D profile of the reflectivity (R), defined as the ratio of neutrons scattered from the interface over the incident intensity of the neutron beam, is measured in specular conditions (*i.e.*, the incident angle of the neutron beam is equal to the reflected angle, denoted as θ) as a function of the momentum transfer vector q normal to the interface (defined as $q = 4\pi \sin\theta / \lambda$, where λ is the wavelength of the neutron beam). The measured reflectivity can be linked to a plane-averaged scattering length density (SLD) profile perpendicular to the interface. SLD measures the coherent scattering cross-section of the molecular species that constitutes each interfacial layer, and is linked to their chemical composition and molecular volume, V by:

$$SLD = \sum_i \; b_i \, n_i$$

where $n_i$ is a number density and $b_i$ is the scattering length density of each molecular species.

### Data acquisition

SNR measurements were performed on FIGARO, a time-of-flight reflectometer, at the Institut Laue-Langevin (Grenoble, France). Two angles of incidence θ (0.62° and 3.8°) and a wavelength resolution of 7% dλ/λ were used yielding a momentum transfer range of $0.007 \leq Q_z \leq 0.25$ Å$^{-1}$ and a residual background reflectivity of R~$10^{-7}$. The raw time-of-flight experimental data at these two angles of incidence were calibrated with respect to the incident wavelength distribution and the efficiency of the detector yielding the resulting R(q) profile using COSMOS [4].

Measurements were performed in HKM buffer (pH 7.1) in different isotopic contrasts, such as $H_2O$, $D_2O$, 8.1% v/v $D_2O$ (denominated air-contrast matched water, ACMW, which has an SLD of zero).

### Data modelling

It was performed by minimizing the difference between the experimental data points and the calculated reflectivity profile. The latter was obtained by a model consisting of multi-layers of constant SLD using the Parratt's recursive method, with an error function connecting adjacent layers. The calculated SLD profiles were convolved by a width 3 Å Gaussian function to approximate the experimental resolution limit. Data analysis was performed using constrains between layer parameters (thickness, roughness and degree of hydration) and simultaneous co-refinement of all data sets obtained with a global minimization of a least-squares function $\chi^2$ was done to reduce the ambiguity in the modelling by using both AuroreNR [5] and Motofit [6] packages.

In the case of lipid monolayers prior AP2 incorporation, a reflectivity profile is calculated based on a real space description of a mixed lipid membrane leaflet. using a two-layer model. An upper layer in contact with the air phase containing the lipid acyl chains and a bottom one close to the bulk phase that includes the lipid polar headgroups. The SLD in both layers is re-parametrized as

$$\rho_i = f_i^w \rho_i^w + f_i \rho_i,$$

Where $f_i^w$ is the volume fraction occupied by the solvent with an SLD $\rho_i^w$ in the layer *i* (≡ lipid tails, head), and $f_i$ is the volume fraction of the molecular entities with an SLD $\rho_i$. By definition $f_i^w + f_i = 1$. In our model, the tail volume fraction ($f_{tail}$) was fixed to unity for simplicity, while the hydrated headgroup layer was fitted considering the solvent volume fraction $f_{tail}^w$ with a fixed thickness of the headgroups estimated by the molecular dimensions. Finally, following, the area-per-molecule of lipidic components was fixed to be the same in each layer $A = \frac{b_i}{SLD_i t_i f_i}$ (with *i* = aliphatic tails or head-groups). In detail, $A_{tails}$ is equal to one corresponding to the hydrated headgroups $A_{head}$ that can be expressed as follows

$$A_{tails} = \frac{b_{tails} N_A}{\rho_{tails} t_{tails}} = \frac{b_{head} N_A}{\rho_{head} f_{head} t_{head}} = A_{head},$$

from which we can establish a correlation between the parameters $f_{head}$ and $t_{tails}$ during the global fit of the SNR data recorded in the different isotopic contrasts. We assume in this simultaneous fitting analysis that the data in different contrasts yield similar interfacial structures.

In presence of AP2, in each layer (*i.e.*, tails and heads for mimicking membrane leaflet) the SLD and the volume fraction *f* follows: $\rho_{layer} = f_{solvent}.\rho_{solvent} + f_{lipid}.\rho_{lipid} + f_{AP2}.\rho_{AP2}$; where $f_{solvent} + f_{lipid} + f_{peptide} = 1$. All the parameters used in the fitting process are tabulated in

Table S5. The headgroup molecular volumes and thicknesses were fixed to values obtained from X-ray diffraction measurements, while the acyl tail region, had a molar averaged SLD value (see **Table S5**).

From the most plausible SLD profiles shown in **Figure S15**, it is possible to calculate the corresponding volume fraction distribution plotted as a function of the distance to the interface for both open and close conformations in **Figure 6A**. In a further step of refinement, theoretical reflectivity profiles, calculated based on the SLD and volume fraction profiles of the best orientations, were fitted against the experimental profiles. A representative comparison is plotted as an inset in **Figure 6A**.

### Low $Q_z$ method SNR

Quantification of the AP2 binding to a lipid monolayer was done exploiting by collecting data at a limited $Q_z$ - range, 0.01 Å$^{-1}$ < $Q_z$ < 0.03 Å$^{-1}$ (λ from 4.5 to 13 Å), which allows a relatively fast acquisition time (≈1 min per curve). A one-layer model of the monolayer (**Figure S20**) was exploited to determine the amount of AP2 bound to the monolayer by using

$$\Gamma_{AP2} = \Gamma_L + \frac{(\mathrm{SLD}\ t)_{fit}}{\Sigma \mathrm{b}_{AP2}},$$

Where $\Gamma_L$ corresponds to the lipid monolayer surface concentration prior the injection of AP2 in the buffer subphase.

### Tables related to SNR experiments

**Table S6.** Scattering length ($\Sigma b$), molecular volume ($V_m$) and scattering length density ($\rho$) used in this work for the different samples studied.

| Sample | Components | $\Sigma b$ (fm) | $V_m$ ($Å^3$) | ρ ( $10^{-6}$ $Å^{-2}$) |
|---|---|---|---|---|
| **PtdIns(4,5)$P_2$** | Acyl chains | –15.99 | 772 | –0.21 |
| | Headgroups | 71.60 | 503 | 1.42 |
| **CD4-PtdIns(4,5)$P_2$** | Acyl chains | –17.39 | 766 | –0.23 |
| | Headgroups/D2O | 91.43 | 600 | 1.52 |
| | Headgroups/NRW | 89.23 | 600 | 1.49 |
| **TGN-PtdIns(4,5)$P_2$** | Acyl chains | –17.39 | 766 | –0.23 |
| | Headgroups/$D_2O$ | 95.07 | 616 | 1.54 |
| | Headgroups/NRW | 92.34 | 616 | 1.50 |
| **CD4**[a,b] | CD4 in $D_2O$ | 1085.33 | 3089 | 3.51 |
| | CD4 in NRW | 580.66 | 3089 | 1.88 |
| **TGN** [a,b] | TGN in $D_2O$ | 1069.81 | 3057 | 3.50 |
| | TGN in NRW | 586.87 | 3057 | 1.92 |
| **AP2**[b] | AP2 in $D_2O$ | 79656 | 262027 | 3.065 |
| | TGN in NRW | 73891 | 262027 | 2.82 |

[a]Included in the lipid mixture at a molar percentage shown in Table SX. [b]Values obtained from ISIS Biomolecular Scattering Length Calculator (http://psldc.isis.rl.ac.uk/Psldc/) assuming a 90% of labile hydrogens.

**Table S7.** Summary of the results obtained from the fitting based on a 2 layers model of Langmuir lipid monolayers.

| **Sample** | **Components** | **t (Å)** | ***$f_w$ (%)*** | ***APM* ($Å^2$)** | **$\Gamma$ (μmol/$m^2$)** |
|---|---|---|---|---|---|
| **PtdIns(4,5)$P_2$** | Acyl chains | 13 ± 1 | 0 | 58.0± 0.4 | 2.8 ± 0.1 |
| | Headgroups | 10 ± 1 | 12 ± 2 | 58.0± 0.4 | 2.8 ± 0.1 |
| **CD4** PtdIns(4,5)$P_2$ | Acyl chains | 12 ± 1 | 0 | 63.1± 0.5 | 2.6 ± 0.1 |
| | Headgroups | 10 ± 1 | 10 ± 2 | 63.1± 0.5 | 2.6 ± 0.1 |
| **TGN** PtdIns(4,5)$P_2$ | Acyl chains | 13 ± 1 | 0 | 59.7± 0.3 | 2.8 ± 0.1 |
| | Headgroups/ | 10 ± 1 | 10 ± 2 | 59.7± 0.3 | 2.8 ± 0.1 |

***Table S8***. Summary of the results obtained from the fitting based on a 5 layers model of Langmuir lipid monolayers after the binding of AP2 molecules.

| **Sample** | **Components** | **t (Å)** | ***Coverage (%)*** | ***$f_w$ (%)*** | ***AP2 vol %*** |
|---|---|---|---|---|---|
| **PtdIns(4,5)$P_2$** | Acyl chains | 14 ± 1 | 100 | 0 | 0 |
| | Headgroups | 10 ± 1 | 88 ± 2 | 6± 1 | 6 ± 1 |
| | AP2 Layer **1** | 21 | - | 90 ± 1 | 10 ± 1 |
| | AP2 Layer **2** | 7 | - | 89 ± 1 | 11 ± 1 |
| | AP2 Layer **3** | 35 | - | 95 ± 1 | 5 ± 1 |
| **CD4 PtdIns(4,5)$P_2$** | Acyl chains | 12 ± 1 | 100 | 0 | 0 |
| | Headgroups | 10 ± 1 | 90 ± 1 | 5 ± 1 | 5 ± 1 |
| | AP2 Layer **1** | 15 ± 2 | - | 95 ± 1 | 5 ± 1 |
| | AP2 Layer **2** | 33 ± 2 | - | 86 ± 1 | 14 ± 1 |
| | AP2 Layer **3** | 36 ± 2 | - | 89 ± 1 | 11 ± 1 |
| **TGN PtdIns(4,5)$P_2$** | Acyl chains | 13 ± 1 | 100 | | 0 |
| | Headgroups/ | 10 ± 1 | 90 ± 1 | 5 ± 1 | 5 ± 1 |
| | AP2 Layer **1** | 14 ± 2 | - | 95 ± 1 | 5 ± 1 |
| | AP2 Layer **2** | 38 ± 2 | - | 67 ± 1 | 33 ± 1 |
| | AP2 Layer **3** | 39 ± 2 | - | 89 ± 1 | 11 ± 1 |

**Table S9.** Summary of the results obtained from the fitting based on a 5 layers model of Langmuir lipid monolayers after the binding of AP2 molecules.

| **DOPC:DOPE: PtdIns(4,5)$P_2$:CD4 bilayer** | t (Å) | $f_w$ (%) | CD4 mol% (*v/v*)* | APM (Å$^2$) | t (Å) | $f_w$ (%) | AP2 % (*v/v*) | APM (Å$^2$) |
|---|---|---|---|---|---|---|---|---|
| (2) Water-layer | 5±1 | 100 | | | 7±1 | 100 | 0 | |
| (3) Headgroups-layer | 10±1 | 17±1 | 5 (40)* | 59±9 | 10±1 | 17±1 | 0 | 59±7 |
| (4) Tails-layer | 17±1 | 1.8±0.2 | | 59±10 | 17±1 | 3.3±0.5 | 0 | 60±4 |
| (5) Tails-layer | 13 | 11.3±0.4 | | 87±10 | 13±1 | 16±1 | 0 | 92±8 |
| (6) Headgroups-layer | 8±1 | 44±1 | 2 (20)* | 88±13 | 8±1 | 29±1 | 20±1 | 90±13 |
| (7) AP2-layer | | | | | 61±2 | 96.2±0.2 | 3.8±0.2 | |

**Figures related to SNR experiments**

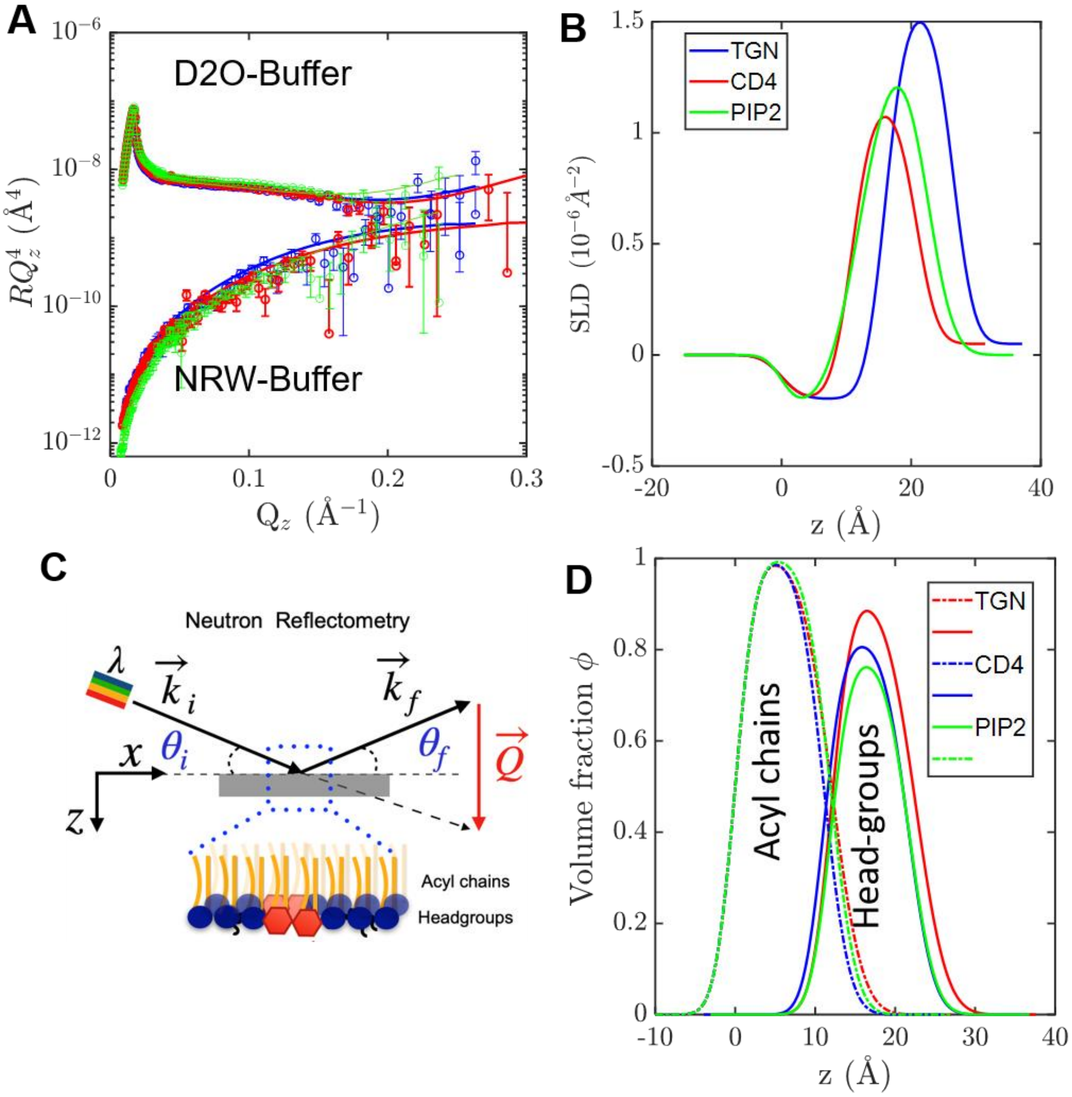

**Figure S10. Specular Neutron Reflectometry on Langmuir lipid monolayers**. **(A)** Neutron reflectivity data of the different hydrogenated lipid monolayers in $D_2O$ and ACMW buffers, respectively. The fitting curves of TGN lipids (blue), CD4 (red) and PIP2 (geen) are also shown. The figure is displayed on a $RQ^4$ scale to show the agreement between the experimental and theoretical data at higher q-values. (**B)** Scattering length density, SLD, profiles corresponding to fits are plotted in **A**. **C** Volume fraction profiles normal to the interface of monolayers to highlight the distribution of acyl chains and headgroups.

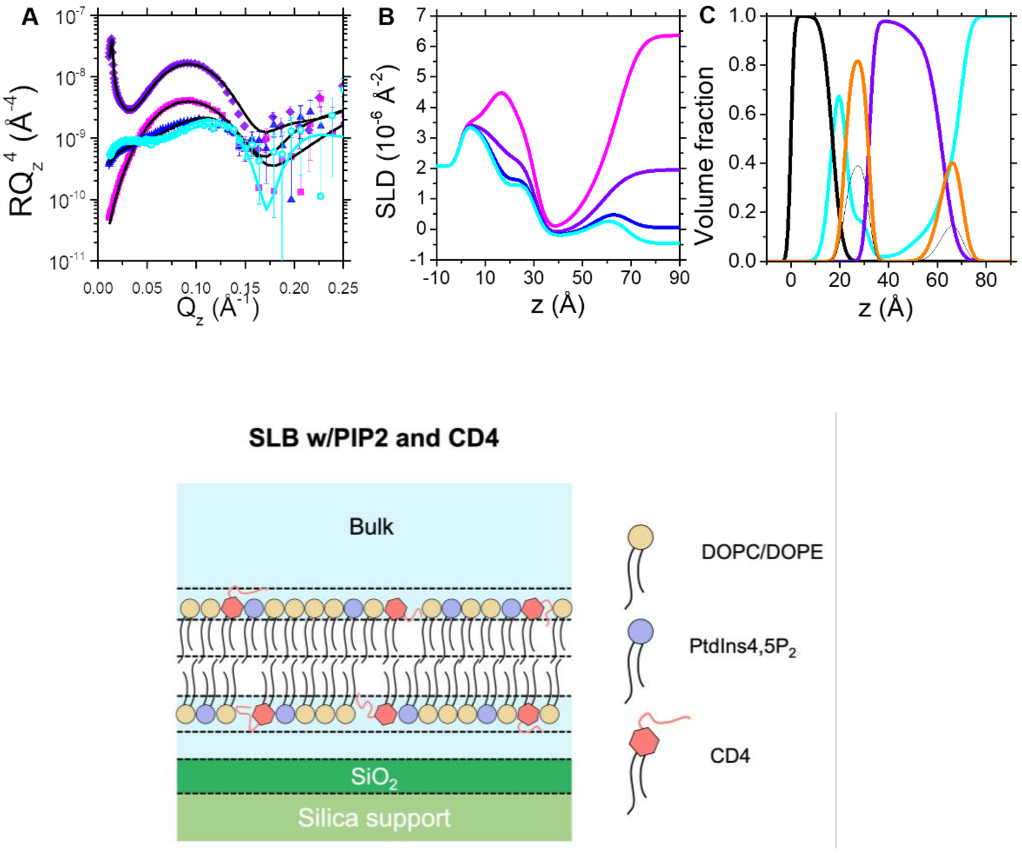

**Figure S11.** (**A**) Reflectivity profiles of DOPC:DOPE:PtdIns4,5$P_2$:CD4 (66.5:20:10:3.5) in 4 contrasts: $H_2O$ (dark blue circle), ACMW (blue triangles), SiMW (violet squares) and $D_2O$ (pink diamonds). The relative fits are presented as lines. (**B**) shows the SLD profiles perpendicular to the interface. (**C**) reports the corresponding volume fraction profiles perpendicular to the interface, showing the contribution of silicon oxide (orange line), aliphatic tails (black line), hydrophilic headgroups (magenta), and water (cyan line). The volume fraction related to CD4 peptide moiety is shown as light pink area.

(D) Schematic representation of the SLB surface after incorporation of PtdIns4,5$P_2$ lipids (shown as purple circles) and in the presence and absence of the lipid-peptide conjugate CD4 (shown as salmon hexagon). The NR fitted parameters for the layers 1 to 6 can be found in ***Table S9***.

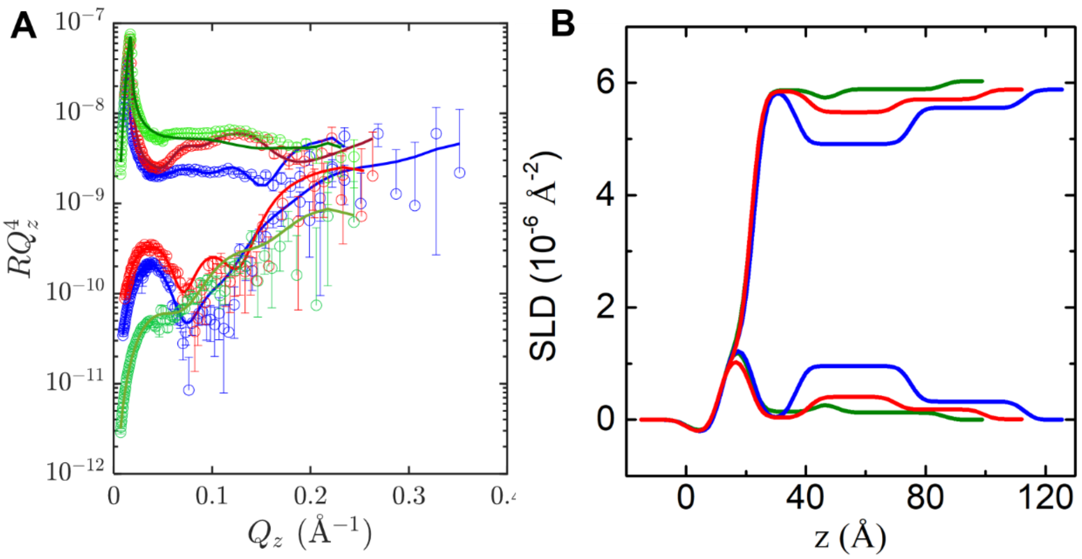

**Figure S12**. Specular Neutron Reflectometry on Langmuir lipid monolayers enriched in lipopeptides after the binding of AP2. Color code as follows: green corresponds to PtdIns(4,5)$P_2$ -monolayer, red to CD4-enriched monolayer and blue to TGN-enriched one. Data are collected in D2O buffer (top data) and ACMW buffer (bottom data).

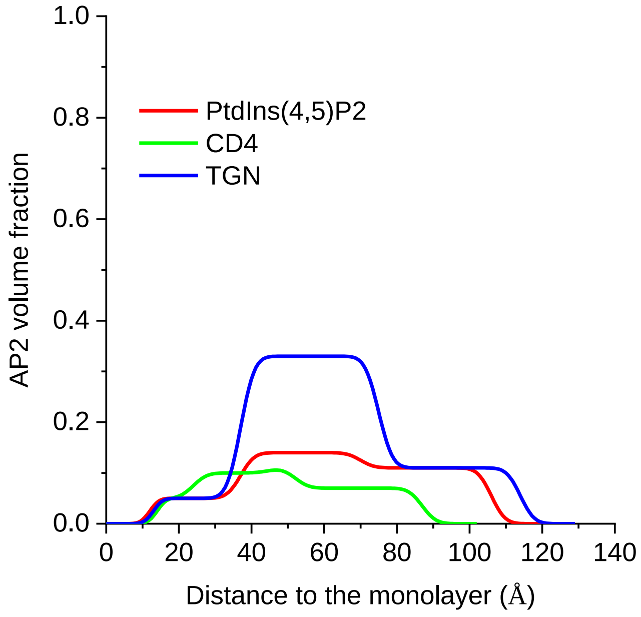

**Figure S13**. AP2 volume fraction profiles plotted against the distance to the lipid monolayer derived from the analysis of the specular neutron reflectometry data showed in Figure S12.

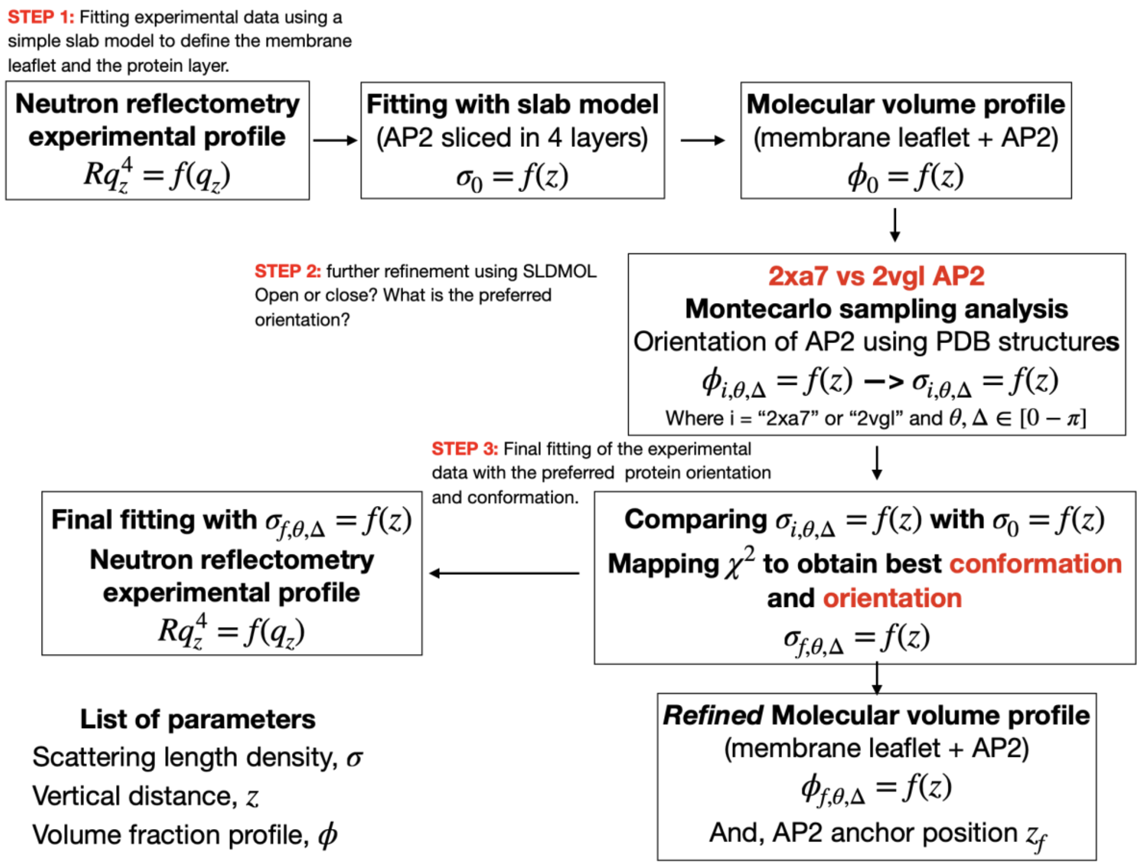

***Figure S14. Block diagram showing the different steps in SNR analysis performed.***

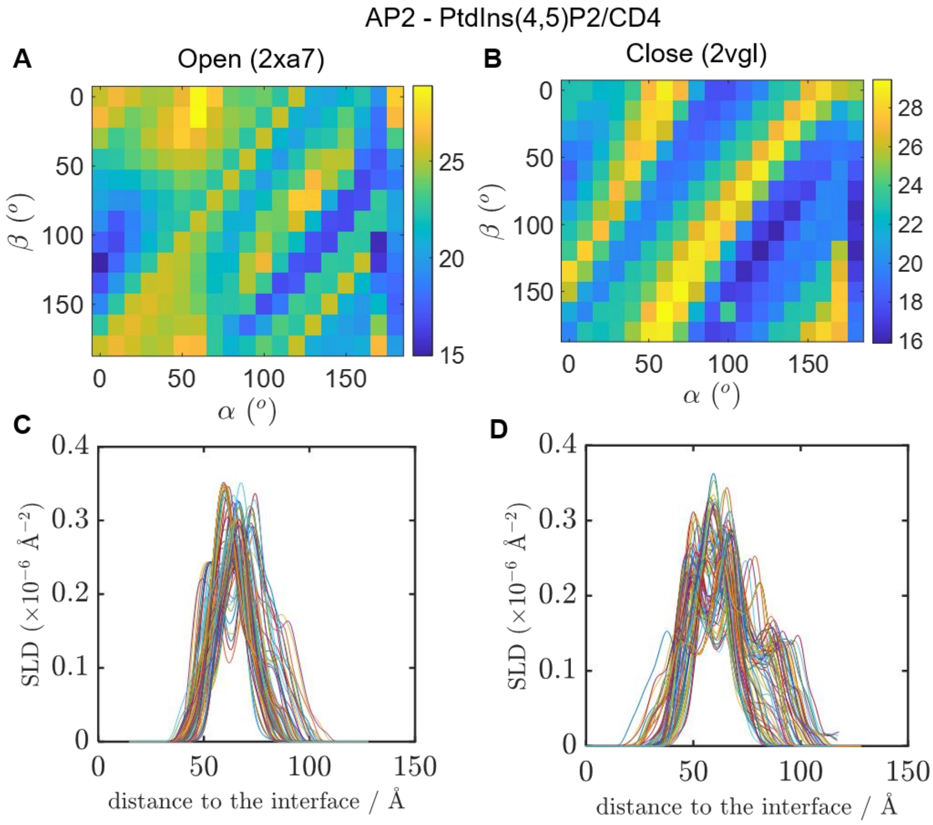

**Figure S15 Atomic structures of AP2 interacting with CD4 enriched monolayers**. Energy minimized crystal structures from PDB 2ax7 and from 2vgl. used to position and orient AP2 component of the experimentally-derived SLD profile. The orientations were parameterized using the Euler angles $\alpha$ and $\beta$ (following an x-y-z extrinsic rotation scheme). An ensemble of 246 different orientation trialled, with 5$^{o}$ increments in $\alpha$ and $\beta$. A and B show colormaps of the preferred orientations for 2xa7 and 2vgl, whilst c and d the respective SLD profiles generated for AP2 as a function of the distance to the interface.

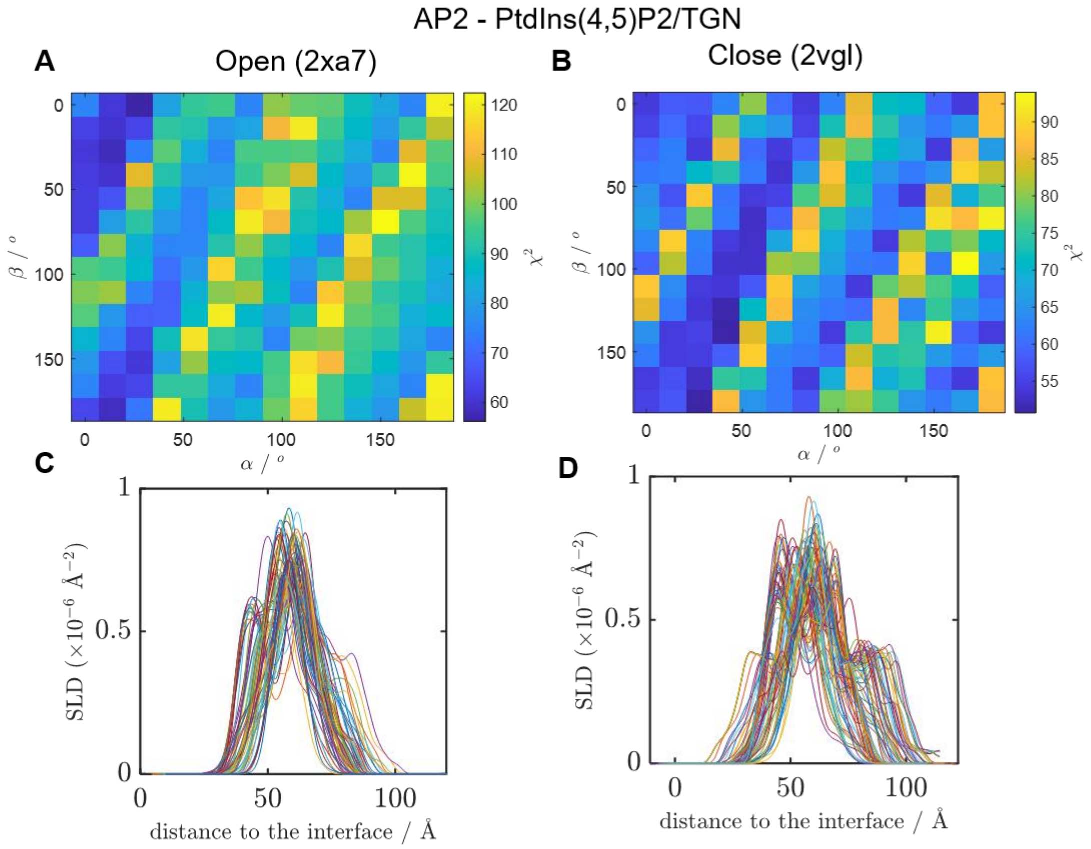

**Figure S16. Atomic structures of AP2 interacting with TGN enriched monolayers**. Energy minimized crystal structures from PDB 2ax7 and from 2vgl. used to position and orient AP2 component of the experimentally-derived SLD profile. The orientations were parameterized using the Euler angles $\alpha$ and $\beta$ (following an x-y-z extrinsic rotation scheme). An ensemble of 246 different orientation trialled, with 5$^{o}$ increments in $\alpha$ and $\beta$. A and B show colormaps of the preferred orientations for 2xa7 and 2vgl, whilst c and d the respective SLD profiles generated for AP2 as a function of the distance to the interface.

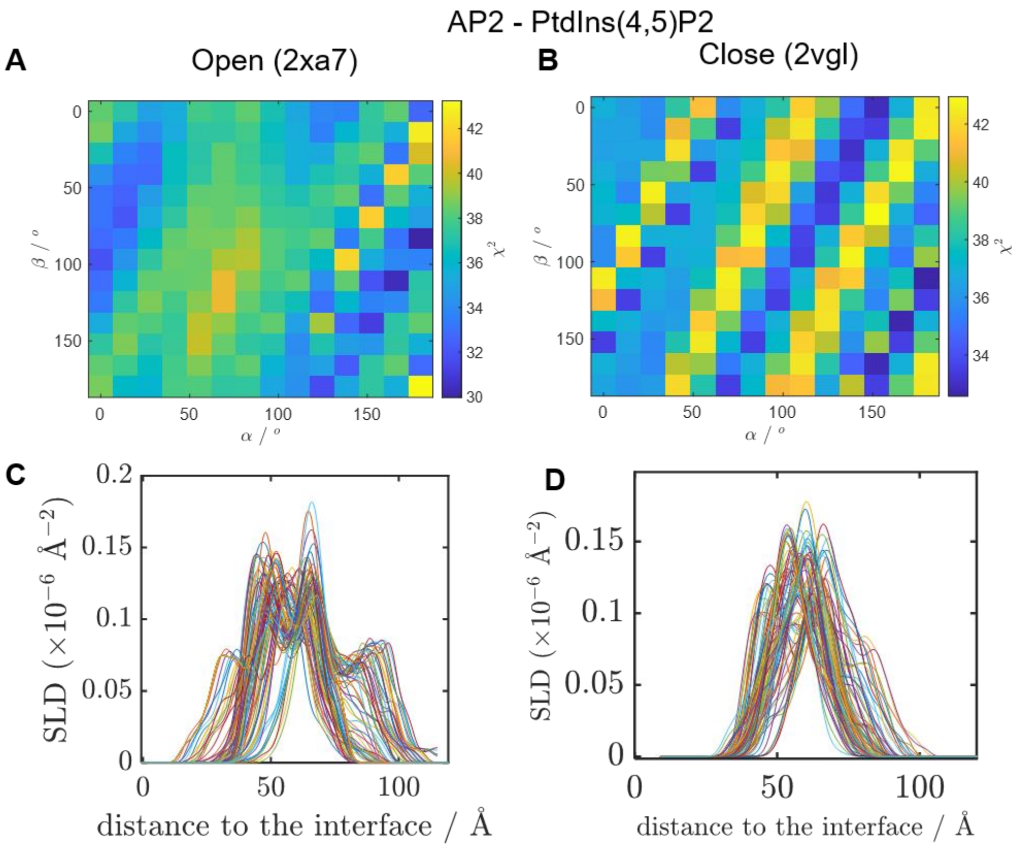

**Figure S17**. **Atomic structures of AP2 interacting with PtdIns(4,5)P2 enriched monolayers**. Energy minimized crystal structures from PDB 2ax7 and from 2vgl. used to position and orient AP2 component of the experimentally-derived SLD profile. The orientations were parameterized using the Euler angles $\alpha$ and $\beta$ (following an x-y-z extrinsic rotation scheme). An ensemble of 246 different orientation trialled, with 5$^{o}$ increments in $\alpha$ and $\beta$. A and B show colormaps of the preferred orientations for 2xa7 and 2vgl, whilst c and d the respective SLD profiles generated for AP2 as a function of the distance to the interface.

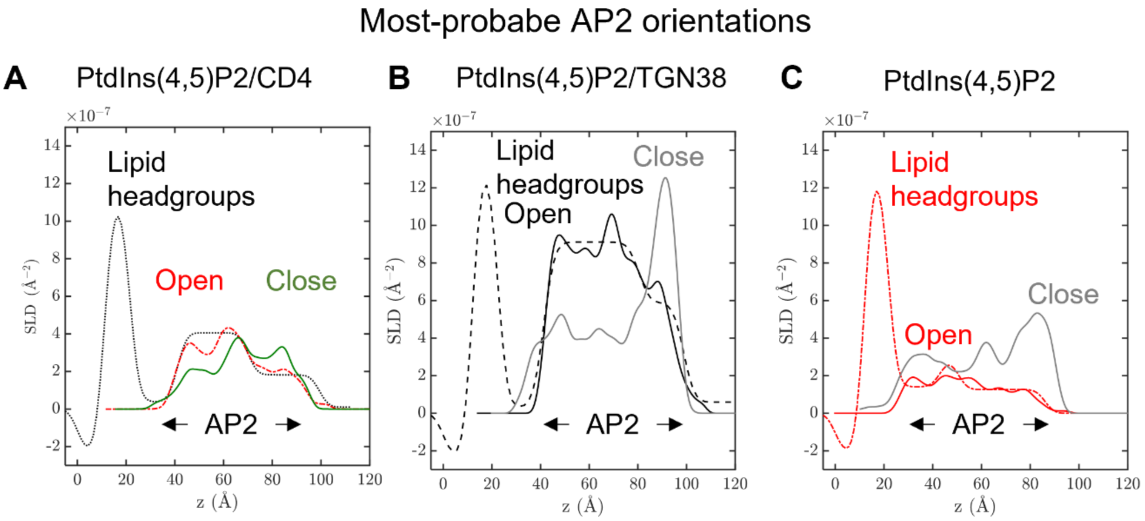

**Figure S18. Most probable AP2 orientations from high resolution SNR.** Comparison of the favored orientations of AP2 in presence of CD4 (A), TGN (B) and PtdIns(4,5)P2 (C) enriched monolayers for the open (2xa7) and close (2vgl) conformations (lowest $\chi^2$) with the experimental scattering length densities (SLDs) obtained by NR.

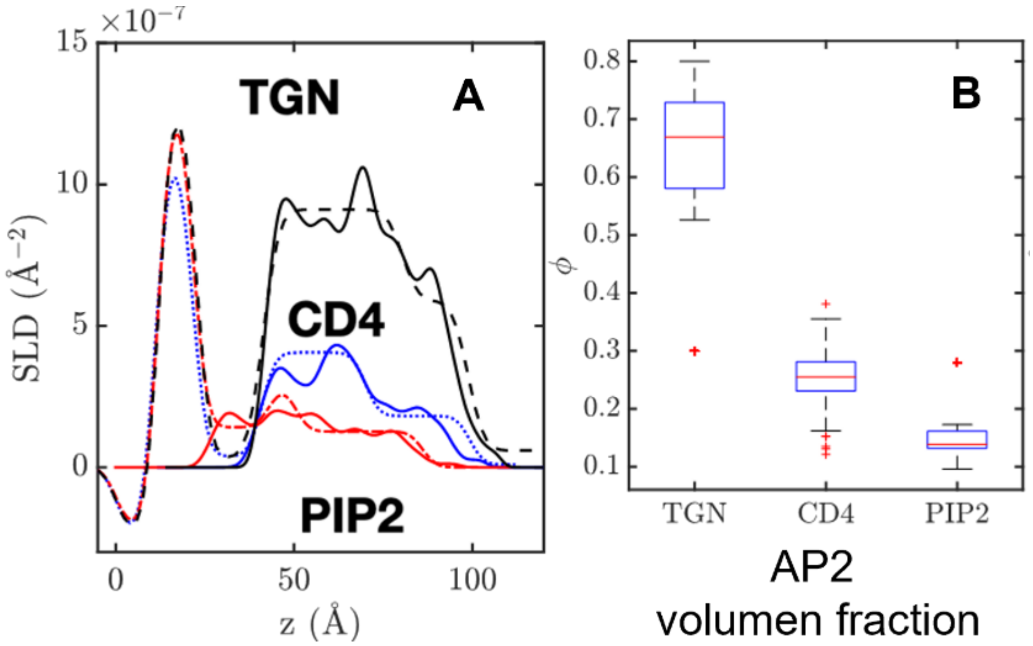

**Figure S19**. (A) Comparison of the best orientation CD4, TGN and PtdIns(4,5)P2 enriched monolayers for open (2xa7) conformation with the experimental SLDs. (B) Corresponding values of volume fractions obtained from the fitting using the theoretical SLD profiles obtained.

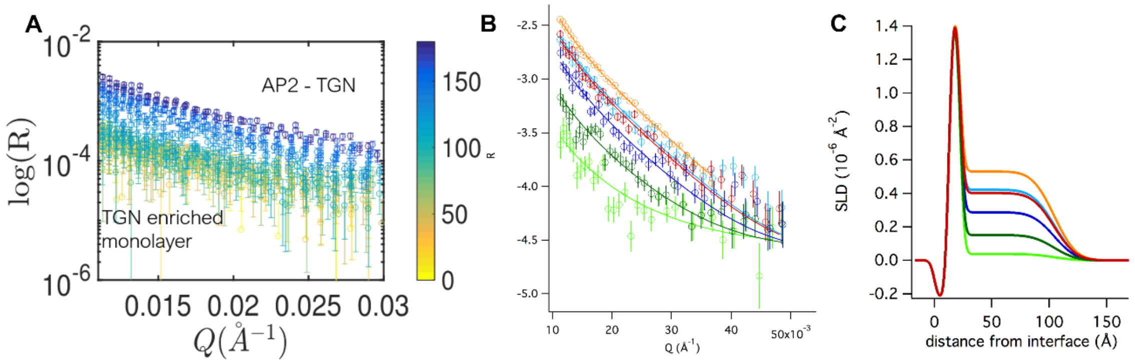

**Figure S20. SNR – low $Q_z$ analysis**. (A) Reflectivity profiles in the region 0.01 Å$^{-1}$ < $Q_z$ < 0.03 Å$^{-1}$. Color bar indicates the evolution of time after AP2 is injected in the subphase. (B) Selected reflectivity profiles fitted with a one-layer model (straight lines). (C) corresponding SLD profiles as a function of the distance to the interface.

### 9. Binding of AP2 onto Solid-supported model PM bilayers

To further explore the association of AP2 with membranes, selected, solid-supported bilayers (SLBs), composed of PtdIns(4,5)P2 enriched bilayers and CD4 cargo were studied. SLBs were prepared by liposome adsorption and fusion into a solid substrate. Small unilamellar vesicles (SUV) were prepared by dissolving DOPC, DOPE, PtdIns(4,5)P2 and CD4 in chloroform, mixing according to desired membrane composition, (i.e. 3:1:1 molar ratio, respectively), dried under gentle Argon flow and placed in vacuum overnight to ensure evaporation of all solvent. The resulting lipid films were rehydrated at room temperature in buffer at up to 1 $mg \cdot mL^{-1}$ lipid concentration, and vortexed to fully suspend vesicles. Immediately before use for supported lipid bilayer (SLB) formation, the suspension was tip sonicated for 5 min at pulses of 1 s on/off to produce a visually clear solution of SUVs. SLB formed from vesicle fusion on the surface.

Subsequent quartz crystal microbalance with dissipation monitoring (QCM-D) experiments (using a commercial Q-Sense E4 instrument, Q-Sense, Biolin Scientific AB, Göteborg, Sweden) indicated stable bilayer formation (**Figure S21**). In comparison with a lipid monolayer, SNR revealed unchanged membrane organization (**Figure S11** and **Table S9**). The bilayer was described incorporating an asymmetrical structure with two inner and outer solvated head groups connected to two lipid leaflet acyl tail layers with different thickness: 13 Å the outer, in contact with the bulk phase and 17 Å the corresponding to the inner leaflet in contact with the solid support. A wetting layer of 5 Å thickness between the silicon crystal and the inner lipid leaflet was also obtained from the SNR analysis. The total bilayer width of 48 Å agrees with QCM data as well as with literature precedent (Fragneto et al., 2018; Pereira et al., 2023). The characterized bilayers had minimum water content in the acyl tail region thus confirming full lipid coverage. Bilayer integrity was furthermore conserved after AP2 binding (**Figure S21, Table S9)**. By assuming that the structure of both the membrane and its bound AP2 are invariant under isotopic substitution, reflectivity data changes on addition of AP2 are compatible with the partition of AP2 into the membrane's outer headgroups region (≈ 20% volume occupancy). The best model form the SNR analysis incorporates an extra layer to account for AP2 present outside the bilayer, with a thickness of 61 ± 1Å and a volume fraction of 3.8 ± 0.2%. These results agree with previous SNR experiments performed on monolayers, in which AP2 only bound to the headgroups and expand to the bulk with a similar volume fraction close to the membrane.

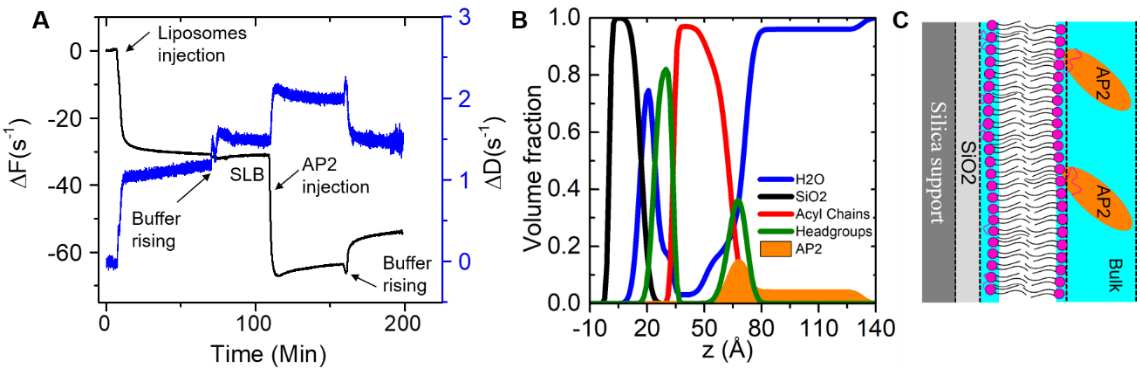

**Figure S 21. SNR and QCM-D data of Lipid bilayers enriched in PtdIns(4,5)P2 and CD4 with AP2**: (A) QCM-D data: Frequency shift and dissipation shift plots (corresponding to the 3rd overtone, n=3) showing lipid vesicles (DOPC, DOPE, PtdIns(4,5)P2 and CD4) adsorption and fusion kinetics: lipid vesicles and subsequent AP2 injection. (B)Volume fraction profiles of solid supported lipid bilayers derived from SNR and SLD profiles plotted in Figure S11. (C) Scheme of the resulting AP2 low-resolution structure bound to lipid bilayers.

### Quartz Crystal Microbalance with dissipation monitoring

QCM-D experiments were performed using a commercial Q-Sense E4 instrument (Q-Sense, Biolin Scientific AB, Göteborg, Sweden) fitted with SiO2-coated AT-cut quartz sensors (QSX 303, Q-Sense, Biolin Scientific AB, Göteborg, Sweden). These sensors were thoroughly cleaned, prior to their use, in an ultrasound bath by sequential immersion in chloroform, acetone, ethanol and water. Afterwards, the cleaned sensors were dried under a gentle stream of nitrogen and exposed to UV-ozone cleaning in a ProCleaner™ Plus instrument (BioForce Nanosciences, Virginia Beach, VI, USA) for 30 minutes. The crystals were then immersed in water and dried under a gentle stream of nitrogen. The cleaned and hydrophilized crystals were immediately fitted inside the QCM-D flow module. The flow module was then connected to a peristaltic pump (Multichannel Peristaltic Pump IPC-N 4, Ismatec, Switzerland) operating with a fixed flow rate of 0.10 mL.min-1.

QCM-D measures the impedance spectra of the quartz crystal for the fundamental frequency ($f$ = 5 MHz) and odd overtones up to the 13th. Before the experiments, the fundamental frequencies of the overtones were recorded up to the equilibration of the signal at the experimental temperature (20 ºC), which is evidenced for a stable baseline, i.e., the $\Delta$f and $\Delta$D remaining constant for at least 5 minutes. After equilibration in buffer, the lipid vesicles dispersion (0.10 mg.mL-1) is introduced in the flow cell and left under incubation for 20-40 minutes, allowing the adsorption and fusion processes. The measurements were performed using freshly prepared vesicles that were stored at a temperature of 4ºC for a maximum of ~ 12 hours (overnight). The QCM-D allows monitoring, simultaneously, the changes of the Δf and ΔD over time. The time evolution of the frequency (Δf/n, where n is the overtone number) and dissipation (ΔD) shifts for the 3rd overtone (n = 3) during the formation of SLBs on the substrate surface from unilamellar lipid vesicles was reported in Figure 5A. Upon injection of the liposomes DF decreases and, simultaneously, the dissipation factor increases until a plateau is reached. Stable Δf value of 30 $s^{-1}$, and ΔD of 1.5 $s^{-1}$ after rising step, indicates stable bilayer formation (Richter & Brisson, 2003, 2005).
